# Heavy-tailed abundance distributions from stochastic Lotka-Volterra models

**DOI:** 10.1101/2021.02.19.431657

**Authors:** Lana Descheemaeker, Jacopo Grilli, Sophie de Buyl

## Abstract

Microbial communities found in nature are composed of many rare species and few abundant ones, as reflected by their heavy-tailed abundance distributions. How a large number of species can coexist in those complex communities and why they are dominated by rare species is still not fully understood. We show how heavy-tailed distributions arise as an emergent property from large communities with many interacting species in population-level models. To do so we rely on stochastic logistic and generalized Lotka-Volterra models for which we introduce a global maximal capacity. This maximal capacity accounts for the fact that communities are limited by available resources and space. In a parallel ‘ad-hoc’ approach, we obtain heavy-tailed abundance distributions from non-interacting models through specific distributions of the parameters. We expect both mechanisms, interactions between many species and specific parameter distributions, to be relevant to explain the observed heavy tails.

---

The abundance distribution of microbial communities is very broad, with many rare taxa and few very abundant ones. These broad distributions are the results of the interplay between interactions and stochasticity in community dynamics. Which ecological mechanisms lead to species abundance distributions is not fully understood. If species abundances were the result of random walks — the process only involving constant noise, for example through immigration and abundance-independent emigration — the central limit theorem would tell us that the abundances follow a normal distribution. Neutral models Hubbell (2001), by assuming random birth and death with species-independent rates, reproduced broad abundance distributions, but fails in explaining stable species differences Grilli (2020) and their correlated patterns. The evolution of a community is not solely governed by random processes, involving interactions between species and the environment.

Most microbial community experimental data fit best lognormal distributions Shoemaker et al. (2017). Those distributions reflect the statistical properties of species temporal fluctuations and long and stable species differences Grilli (2020). Here we explore how lognormal abundance distributions can arise as an emergent property from large communities with many interacting species.

Heavy-tailed distributions can result from self-organized critical behavior (Bak et al., 1987) in individual-based models (IBMs)Solé et al. (2002), as the product of interactions between individuals and immigration of individuals. All individuals of a species are assumed to be equivalent, *i.e*. differences between individuals of the same species are not taken into account: they all have the same growth rates, self-interaction strengths and probabilities of immigration. Species abundance evolve stochastically, where the per-capita birth and death rates also depend on other species’ abundances. We propose here an explicit implementation of IBMs in a generalized framework, and provide parameter values leading to abundance distributions compatible with those of observed microbial communities.

Inspired by IBMs, we propose here population-level models (PLMs) that are also able to reproduce the observed abundance distributions. More specifically, the models we propose are stochastic generalized Lotka-Volterra equations (sgLV) with a global maximal capacity mimicking the finite resource availability (e.g., the finite size lattice on which IBMs are constructed). Those models exhibit heavy tails as an emergent property, originating from the interactions between species, including the effective interactions originating from the maximal capacity. Additionally, we find a power law relation for high connectance values but the diversity *S* saturates for small connectance because there is a finite number of species and a maximal number of individuals in the model. This conclusion is in agreement with Solé et al. (2002) and we explored how it translates to PLMs. We refer to Supporting results for results about this complexity-stability relationship.

An heavy tailed abundance distribution is not the only characteristic exhibited by microbial communities. We previously extensively studied stochastic properties of microbial communities time series and showed that stochastic logistic models could reproduce all those properties in Descheemaeker and de Buyl (2020). We can extrapolate easily those results to sgLV models with maximal capacities. Moreover the maximal capacity proposed nicely prevents unstable solutions.

This paper is organized as follows. We first present an analysis of experimental data of microbial communities and propose different observables to characterize the abundance distribution and time series. Not only do we characterize the width of the abundance distribution but also its temporal fluctuations as well as the coefficient of variation of the fluctuations of individual species. Next, we reproduce and generalize the results of Solé et al. (2002): heavy-tailed distributions can be obtained by IBMs. We study the conditions under which those IBMs reproduce not only lognormal abundance distributions but also the characteristics just mentioned. Afterward, we show, inspired by the set-up of the IBM, how heavy-tailed distributions can also be obtained by stochastic logistic and sgLV models by including a maximal capacity. We next approach the problem from another, simpler perspective. We consider the logistic model without a maximal capacity and show that even without complex interactions heavy-tailed abundance distribution can be obtained by choosing parameters wisely. More specifically, we show that non-interacting models lead to lognormal abundance distributions when the parameters follow lognormal distributions and that power law distributions can be obtained by uniformly distributed parameters.

## Results

### Abundance distributions of experimental data is lognormal

We use experimental data of plankton and microbial communities related to the human body. The sources of all datasets can be found in Sources experimental data David et al. (2014); Martin-Platero et al. (2018); Caporaso et al. (2011); Arumugam et al. (2011); Qin et al. (2010); Turnbaugh et al. (2009). Figure 1A shows that he abundance distributions of the microbial communities under consideration are heavy-tailed and the best fit is obtained by a lognormal function rather than a power law function (see also Figure 1 in Supplementary figures and Heavy-tailed distributions), consistently to what previously reprted Shoemaker et al. (2017).

**Figure 1.**
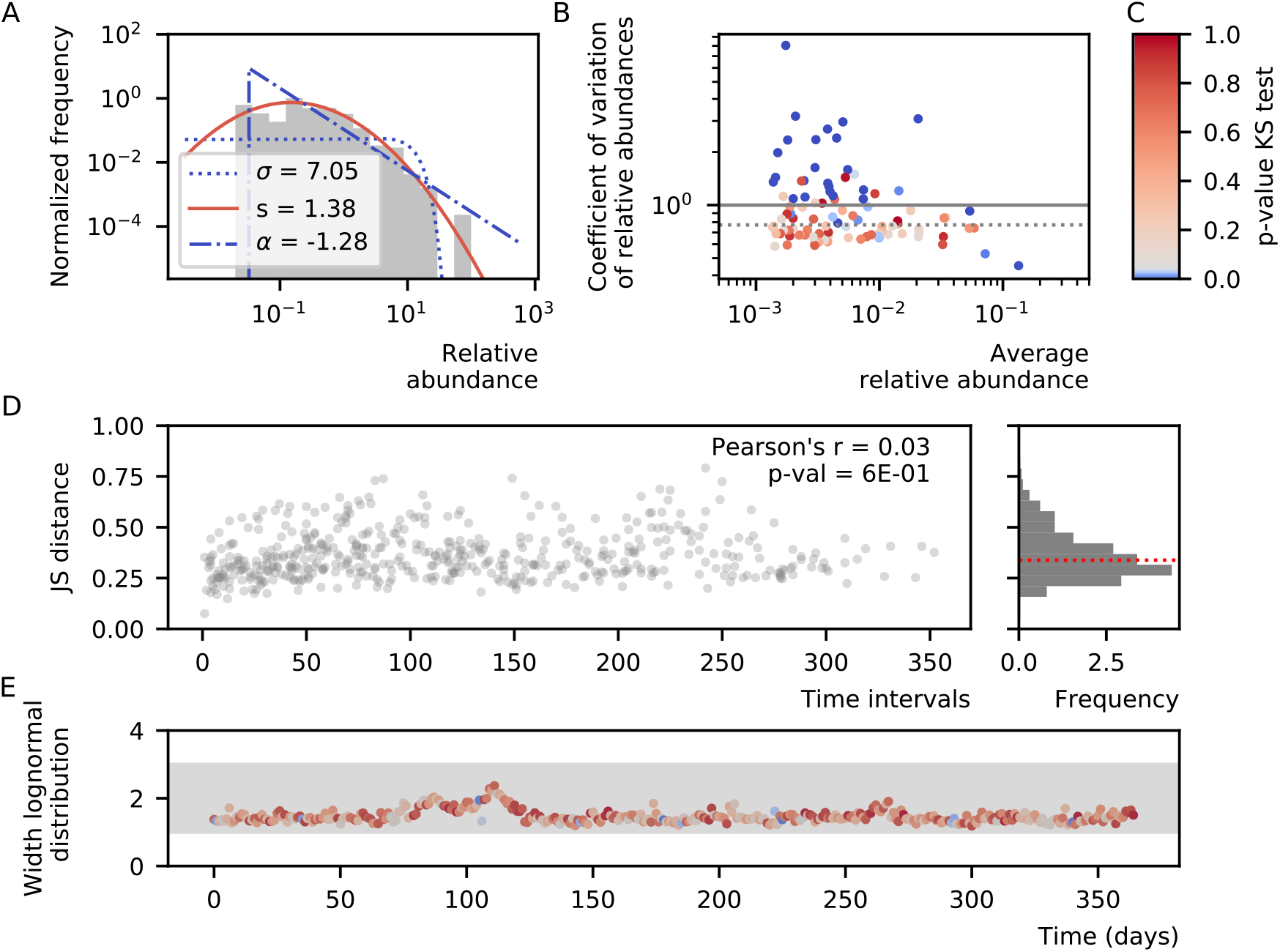
Properties of experimental data. We selected one time series for illustrative purposes and refer to Supplementary figures for time series listed in Sources experimental data. A. Experimental data has a heavy-tailed distribution. The lognormal distribution (full line) fits best. The power law (dash-dotted) and exponential (dotted) distribution are rejected for most communities because the p-values of the KS test are lower than 0.05 (lines are colored in blue). The widths *s* of the lognormal distribution range between 1.42 and 2.78. For data coming from a time series, we considered the first time point. B.Coefflcient of variation of longitudinal experimental data – Longitudinal experimental data of microbial communities has a coefficient of variation that is around one. For species with a large coefficient of variation, we can reject the hypothesis that the population follows a Gamma distribution. The dotted horizontal lines denote the median coefficient of variation per community. C. The color code for the KS test for A., B. and E. D. Jensen-Shannon distance of longitudinal experimental data – Jensen-Shannon (JS) distance of longitudinal experimental data as a function of the time interval between both compositions. The median value (red line) ranges from 0.3 to 0.4. The Pearson correlation function and p-value for non-correlation are given. E. Stability of heavy-tailed distribution of experimental data – The species abundances of microbial communities show large fluctuations over time, but the width of the abundance distribution remains nearly stable over time. The grey horizontal band extends from 1 to 3. The color of the dots shows the p-value of the KS test to a lognormal distributions, blue values are poor fits. The lognormal fits for which the width is higher than 3, can all be rejected.

In Figure 1A and Figure 1 in Supplementary figures, we show the fit of a normal (dotted line), lognormal (solid line) and power law distribution (dot-dashed line). The color of the line represents the p-value of the Kolmogorov-Smirnov(KS) test (details about this test can be found in Kolmogorov-Smirnov test). This means that higher values (darker red) correspond to a better fit, and that fits with values smaller than 0.05 can be rejected (blue). In particular, the normal distribution is rejected for all considered microbial communities and the power law distribution for most of them. In general, a lognormal distribution fits the data best. In the legend, the most important parameter of the different distributions are given, *i.e*. for the normal distribution, the standard deviation *σ*, for the lognormal distribution, the width *s* as defined in Equation 1 and for the power law, the exponent *α*. The lognormal distribution can be described by

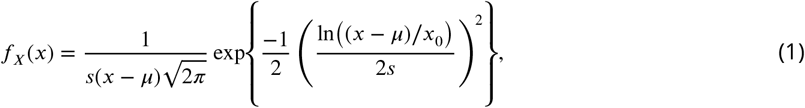

where *μ* defines the location of the distribution, *x*_0_ the scale, and *s* a measure for the width of the distribution. In experimental data, the latter ranges from 1.42 to 2.78 (Figure 1A and Figure 1 in Supplementary figures). Although the species abundances show large fluctuations over time, the width of the abundance distribution remains nearly stable (Figure 1E and Figure 2 in Supplementary figures)

An alternative for studying the abundances of communities is the rank abundance curve. For power law abundances, the rank abundance is also a power law (see Rank abundance distribution). For the lognormal abundance distribution, the width *s* reflects the steepness of the rank abundance distribution. Larger values of the width *s* result in steeper curves (Figure 2).

**Figure 2.**
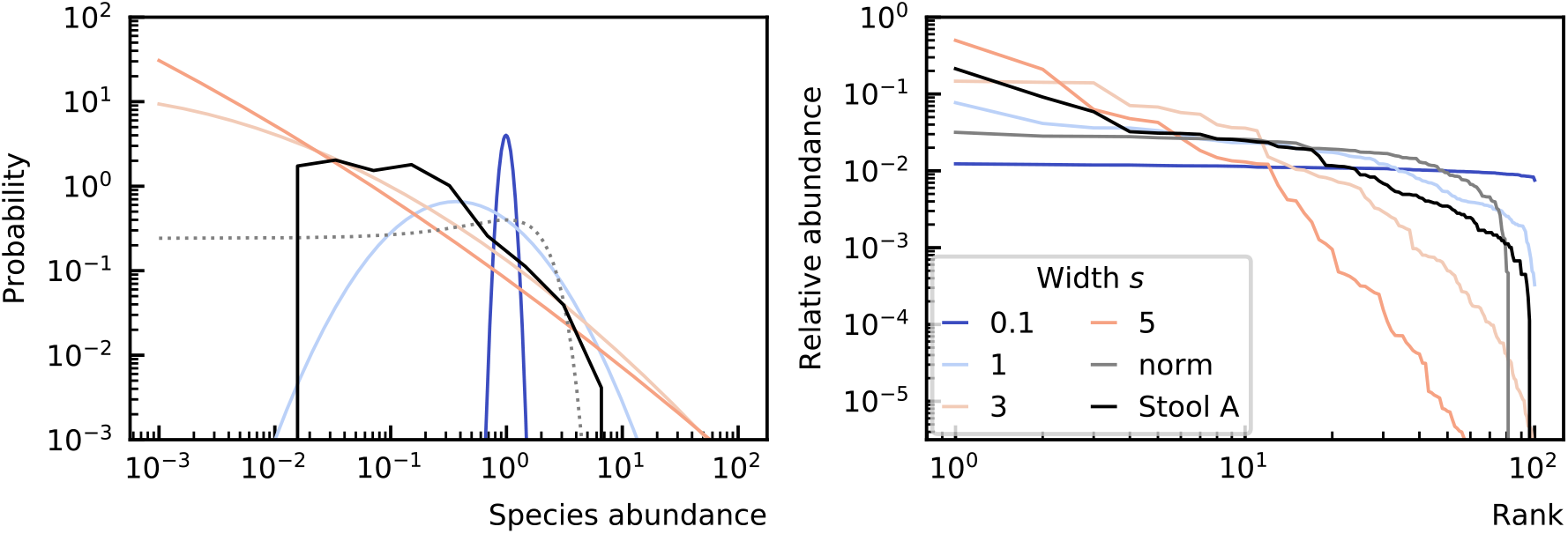
The width of the lognormal distribution *s* reflects the steepness of the rank abundance curve. The larger the width of the abundance distribution, the steeper the rank abundance curve. For instance the community represented in the darker blue has the smallest width for its species abundance distribution and the flatter rank abundance curve. The opposite case is represented by the communities in red exhibiting largest width for their species abundance distribution and steepest rank abundance curves. ‘Stool A’ refers to the time series Stool A of David et al. (2014).

However, it is not sufficient to find models that lead to heavy-tailed distributions. We want the temporal fluctuations of the modeled time series not to be larger than fluctuations in experimental data. We can characterize the fluctuations of the composition as a whole by considering the Jensen-Shannon (JS) distance between microbial compositions at different time points. The definition of the JS distance can be found in Dissimilarity measures. Basically, it is a metric that is used to describe the distance between community compositions. The median JS distance between states at different time points in experimental data ranges from 0.3 to 0.4 (Figure 1D and Figure 4 in Supplementary figures). This is for longitudinal data (multiple time points of one community). For comparison, in cross-sectional data of the gut (different communities at one time point) the median distance ranges from 0.4 to 0.5 (Supporting results). We calculated the Pearson correlation coefficient for the JS distance as a function of the length of the time interval between the two abundance distribution considered and the p-value for the hypothesis of non-correlation. For the planktonic eukaryotes, we reject the hypothesis that there is no correlation between the time interval and JS distance. The community is not stable and is slowly evolving which can be explained by a seasonal change in the seawater (Martin-Platero et al., 2018). On the other hand, for the stool data of subject A, this hypothesis cannot be rejected indicating that the community fluctuates around a steady state as we expected.

Another way of assessing the strength of the fluctuations is by considering fluctuations of separate species. According to Grilli (2020), the distribution of relative species abundance at different time points is a Gamma distribution. The coefficient of variation — the ratio of the standard deviation to the mean — is a variable that determines this distribution (see Heavy-tailed distributions). We find that some of the species indeed follow a Gamma distribution, specially when the coefficient of variation is small. In Figure 1B and Figure 3 in Supplementary figures, the coefficient of variation of all species is shown and the color of the dots represents the p-value of the KS test for the Gamma distribution. This means that we can reject the hypothesis that the abundance distribution over time for a species follows a Gamma distribution for the species with blue points and that the species with dark red points are likely to follow a Gamma distribution. The median coefficient of variation is between 0.7 and 2. We expect the median coefficient of variation to correlate with the median JS distance, as both measures reflect the width of the abundance distribution. However this is not necessarily true as one can imagine a community where all species grow and decrease maintaining the same relative abundances. In this case, the JS distances would remain zero while the coefficients of variation do not.

To sum up we want our models to mimic the characteristics of the experimental data which can be summarized as: time series fluctuate around a steady state with a median JS distance of 0.3 to 0.4, a coefficient of variation between 0.7 and 2 and a lognormal abundance distribution with a width *s* between 1 and 3.

### Revisited Individual Based Models lead to lognormal abundance distributions for low immigration rates and high interaction rates

As mentioned above, Solé et al. (2002) reported that heavy-tailed distributions can be obtained with an IBM with a fixed maximum number of individuals. They argue that ecological systems might be organized at a critical point, balancing immigration of new species with interaction between species. We rely here on the IBM proposed in Heyvaert (2017) which is an explicit interpretation and generalization of the model of Solé et al. (2002) that allows for all types of interactions, *i.e*. not only predation, but also mutualism, competition, amensalism and commensalism. Those IBMs are stochastic cellular automatons in which immigration, growth, and extinction events can happen at every time step as well as the interaction between species. They exhibit heavy-tailed distributions even when all species have equal growth rates, self-interaction, and immigration probability. Every species unevenness is, therefore, a result of stochastic fluctuations or the interaction between individuals of distinct species. Heyvaert(2017) and Solé et al. (2002) show that depending on the immigration rate, both power law and lognormal abundance distributions can be obtained. More details about the exact implementation of this model can be found in Individual-based models.

Before turning to PLMs, we first derive the criteria which need to be met for IBMs to exhibit heavy tails. In this perspective we repeat the analyses of Solé et al. (2002) and Heyvaert (2017) and extend the results in several ways.

We study the influence of the immigration rate as done previously, and additionally study the influence of the connectance of the interaction matrix. The interaction matrix is characterized by two parameters: the interaction strength and the connectance. The interaction strength *a* determines the distribution from which the interaction parameters are drawn: we use the uniform distribution 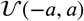. The connectance *C* represents the ratio of interactions (off-diagonal elements of the interaction matrix) that are non-zero. In practice, given the interaction strength *a* and connectance *C*, we first draw all elements of the interaction matrix from the uniform distribution 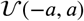 and subsequently put random elements to zero with a probability 1 − *C*. This is similar to the construction of an Erdős-Rényi graph. We impose all remaining parameters to be equal for the different species: the self-interactions *ω_ii_* = −1, the growth and death or emigration rates 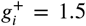 and 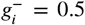 and immigration rate *λ_i_* = *λ*. For various immigration rates and values of the connectance, we did the following: (a) We quantify which distribution fits the abundance distribution best by using the KS test. More details about the fit methods can be found in Fit distributions. (b) Furthermore, when the lognormal distribution has the best fit and is not rejected (p-value of KS test higher than 0.05), we calculate the width of the lognormal distribution. (c) We also computed the average JS distance between different time points, and the average coefficient of variation of the relative abundances of species. Notice that all these measures are defined by the relative abundances of species and not the absolute abundances. The width of the lognormal distribution is independent of the scale, the JS distance is defined for fractional abundances and we choose to measure the coefficient of variance of the relative abundances. As many experimental data is measured as fractional abundances, this facilitates comparing theoretical with experimental results.

We conclude that the abundance distribution shifts from a lognormal to a power law distribution for a decreasing immigration rate and increasing connectance (Figure 3A). This result is in agreement with the results of Solé et al. (2002) with respect to varying immigration rates. The transition from lognormal to power law distributions is gradual with increasing width of the lognormal distribution (Figure 3B). Also, as expected, the number of species is higher when there is immigration but low without immigration because species that go extinct cannot reappear in the community. Moreover both measures of fluctuation — the average coefficient of variation (Figure 3C) and average JS distance (Figure 3D) — increase for decreasing immigration. These results are summarized in Figure 3. They have been obtained for a maximum number of individuals, *N*_max_ = 10^4^. For smaller maximum numbers of individuals, the diversity decreases and the width of the lognormal abundance distribution and the measures of fluctuation increase. This is discussed in Supporting results.

**Figure 3.**
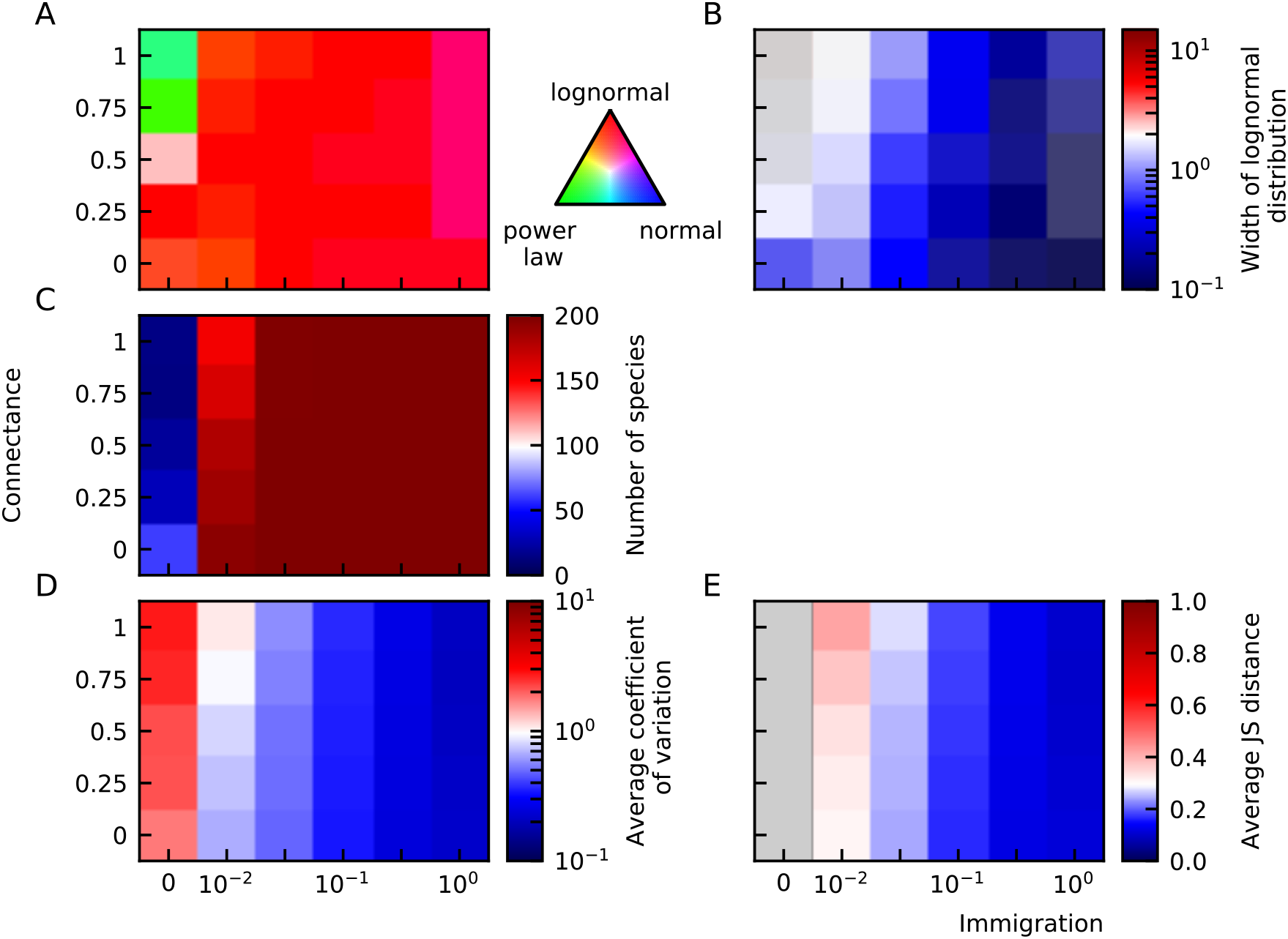
Individual-based models result in heavy-tailed abundance distributions. (A) Abundances mostly fit a lognormal distribution best, with a tendency to power law distributions for small immigration and high connectance. (B) The width of the lognormal distribution increases for decreasing immigration and increasing connectance. (C) The species diversity is maintained except for small immigration rates. The average coefficient of variation (D) and average JS distance (E), increase for decreasing immigration. The fixed parameters of these plots are the number of species *N*_spec_ = 200, the interaction strength *a* = 0.75 and the maximum number of individuals *N*_max_ = 10000. Note that the color code has been chosen such that the white range corresponds to experimental values.

### Heavy-tailed distributions as emergent property of stochastic Lotka-Volterra models with a maximal capacity

We have shown that IBMs can result in heavy-tailed abundance distributions. The main question addressed in this work is whether one obtains qualitatively similar behavior for PLMs. We study the widely used generalized Lotka-Volterra (gLV) equations. For each species *i*, there is an immigration term *λ_i_*, a growth rate *g_i_*, and the self-interaction term *ω_ii_*. This model also includes pairwise interactions between the different species: *ω_ij_* represents the effect of species *j* on species *i* which can be beneficial (*ω_ij_* > 0), harmful (*ω_ij_* < 0) or non-existent (*ω_ij_* = 0). We will consider a linear noise strength *η_i_* as argued in Descheemaeker and de Buyl (2020). The stochastic gLV (sgLV) equation of species *i* reads

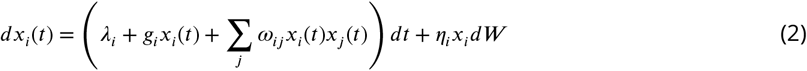

with *dW* an infinitesimal element of a Brownian motion which is defined by a variance of 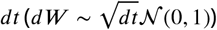. The values for all these parameters were discussed in Revisited Individual Based Models lead to lognormal abundance distributions for low immigration rates and high interaction rates.

The IBM restricts solutions that go to infinity by construction: there is a maximum number of individuals. In gLV models, there is no such restriction and high interaction rates lead to a loss of diversity (blue region in Figure 4A for, *N*_max_ = ∞). Therefore, we do not find the same qualitative behavior as for IBM. To still obtain heavy-tailed distributions with gLV, we want to mimic the IBM. Therefore we compared in details IBM and sgLV models.

**Figure 4.**
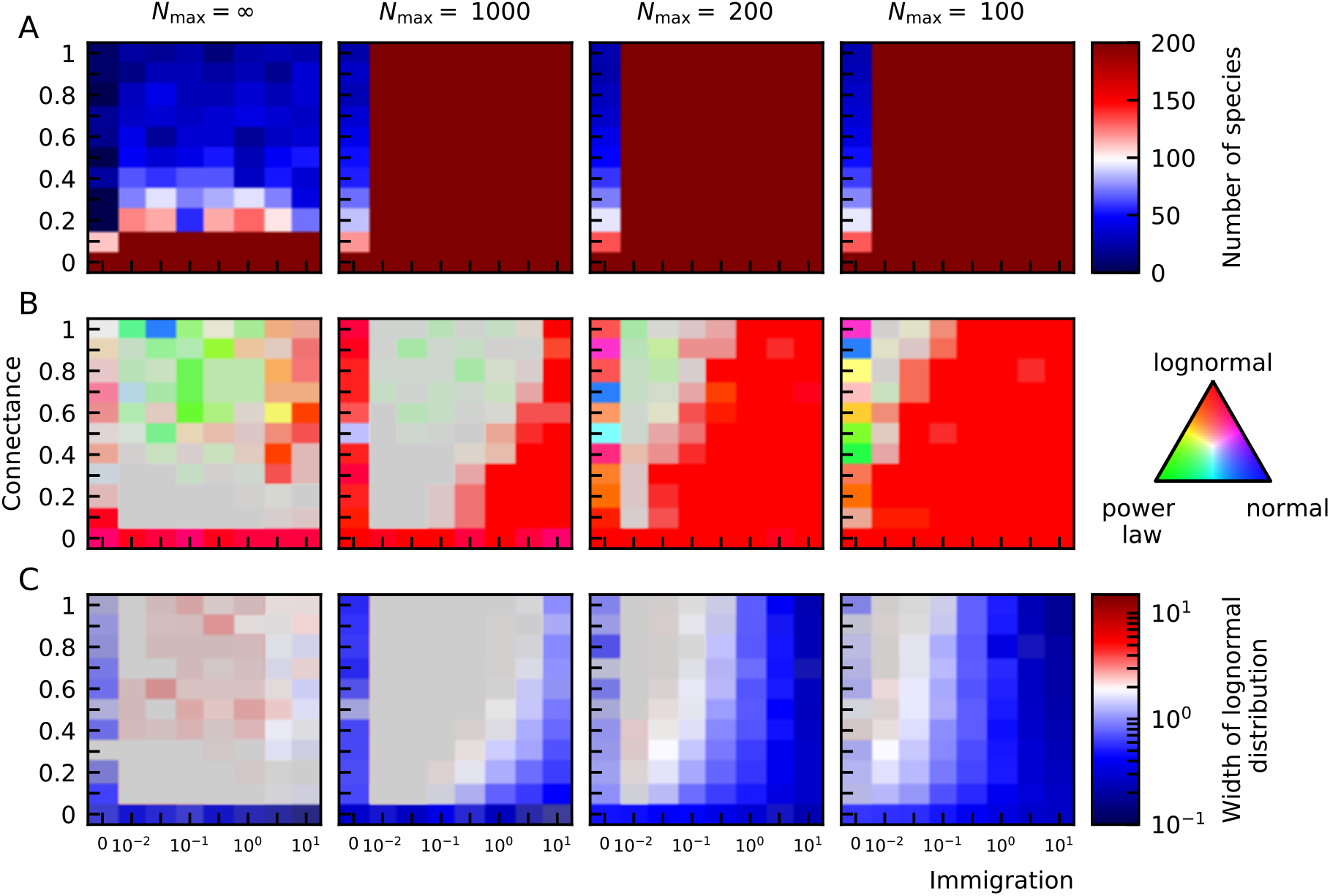
A finite maximal capacity and non-zero immigration rate allow for high diversity in the presence of interactions (A). Notice that the maximal capacity does not limit the number of species, *e.g*. there can be 200 species even when *N*_max_ = 100 because the species abundances can be smaller than 1 in population-level models. For decreasing maximal capacities the abundance distribution becomes lognormal (B). The grey area represents distributions that are neither lognormal, power law or normal. The width of this lognormal distribution increases for decreasing immigration and increasing connectance (C).

For gLV models, one subtlety regarding the extinct species needs to be taken into account. In IBMs species can be absent, in which case they are not considered for the abundance distribution. In gLV models without noise, species that go extinct will take an infinite amount of time to reach zero. Noise or finite integration steps can lead a population to obtain zero or negative abundances (in the latter case, the abundance is put to zero), but otherwise, the species have a very small abundance at the end of the time series. We will consider all species that have an abundance that is six or more orders of magnitude smaller than the maximum abundance in the community as if they were extinct. For the IBMs we performed simulations for a maximum number of individuals up to 10,000 which leads to a difference of maximally four orders of magnitude between populations. This constraint is important for fitting the heavy-tailed distributions, but even more so in the study of diversity-complexity.

The simulation rules of the IBM are based on the gLV model which describes the abundances at the population level. The immigration, extinction, and interaction rates can be chosen to correspond to the parameters of a gLV model, but one important difference is the fixed maximum number of individuals. The growth rate of species, therefore, depends on the number of empty sites, whereas the growth rate in the gLV model is a scalar. Biologically, the maximal capacity can be interpreted as limits by fundamental resources such as water, nutrients, and space. This is why we propose a PLM with a maximal capacity. We implement the maximal capacity to logistic and sgLV models as follows: (1) We multiplying all positive growth terms with a growth probability *γ* that depends on the total number of individuals *N_max_, γ* = 1 − ∑_*j*_ *x_j_*/*N*_max_; (2)The growth vector *g* can be split into its positive and negative components *g* = *g^+^* + *g^-^*. Species for which the net growth term is positive can grow in the absence of other species, species with a strictly negative growth term will need other species to interact positively with them to have a non-zero positive steady-state abundance; (3) Analogously, the interaction matrix can be split into its positive and negative parts: *ω* = *ω^+^* + *ω^-^*. Only the positive growth terms and immigration rate are multiplied by the probability for growth *γ* that depends on the fraction of “empty space”. Furthermore, we limit the probability for growth *γ* such that it has a value between 0 and 1. This constraint is important because we could imagine a steady state starting with more individuals than the maximal capacity. Without a limit at 0, *γ* could become negative and the growth term becomes a death term. The implementation of the maximal capacity *N*_max_ into gLV models (Equation 2) thus resumes:

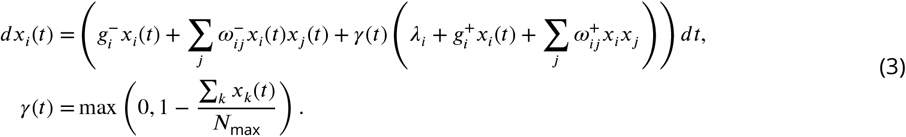

Noise can be implemented by adding a linear random fluctuation *η,x,dW* to the right-hand side of Equation 3 similar to the noise in Equation 2. Notice that the total number of individuals can exceed the total capacity because noise term is not bounded by the maximal capacity. The maximal capacity is, therefore, not a rigid constraint. One could also implement the maximal capacity effect for both the deterministic and stochastic parts, *i.e*. multiply the noise term by *γ*. This gives a bias to negative values for the noise term.

We consider first the simplest case of our model: without interactions, *i.e*. the logistic equations.

#### Logistic equations with a maximal capacity are generalized Lotka-Volterra equations

The logistic equations are similar to the gLV equations (Equation 2), but the immigration and interaction terms are omitted:

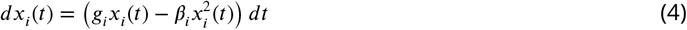

with *β_i_* the self-interaction of species *i*. We do not include the noise term in the next derivations, but one can easily add it to these equations. The self-interaction is related to the carrying capacity *K_i_* = *g_i_*/*β_i_*. After adding a maximal capacity to the logistic equations

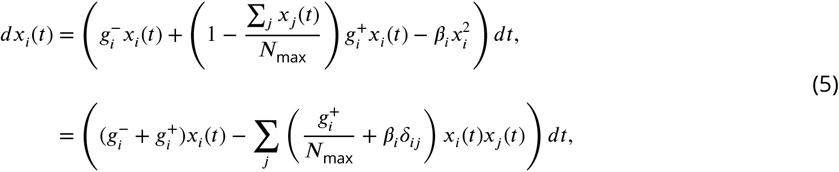

we can identify an interaction matrix 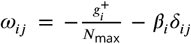. Logistic equations with a maximal capacity can thus be interpreted as gLV equations as long as we assume that the self-interaction has only a negative component 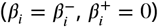. The reduction to a gLV model is valid as long as the sum of the abundances is smaller than the maximal capacity. One needs to be careful to not use initial conditions where ∑_*j*_ *x_j_*(0) > *N*_max_. A smaller maximal capacity translates into more competitive interactions in the gLV model.

Logistic equations with a maximal capacity have thus effective interactions between all species. However, in this work, we use the terminology of interactions only for the specific interactions and not the indirect interactions through the maximal capacity.

The width of the lognormal distribution is determined by the maximal capacity and noise level. We perform simulations for the system of logistic equations with maximal capacity (all growth rates equal to 1, all self-interactions equal to −1, no specific interactions). Both increasing the noise and decreasing the maximal capacity make the lognormal distribution wider (Figure 5). To obtain a width of around 2 (white values), we need a balanced combination of maximal capacity and noise. The smaller the maximal capacity, the less noise is needed to obtain a similar width. We see that decreasing the maximal capacity increases the JS distance (Figure 5D). As expected increasing the noise also increases the JS distance.

**Figure 5.**
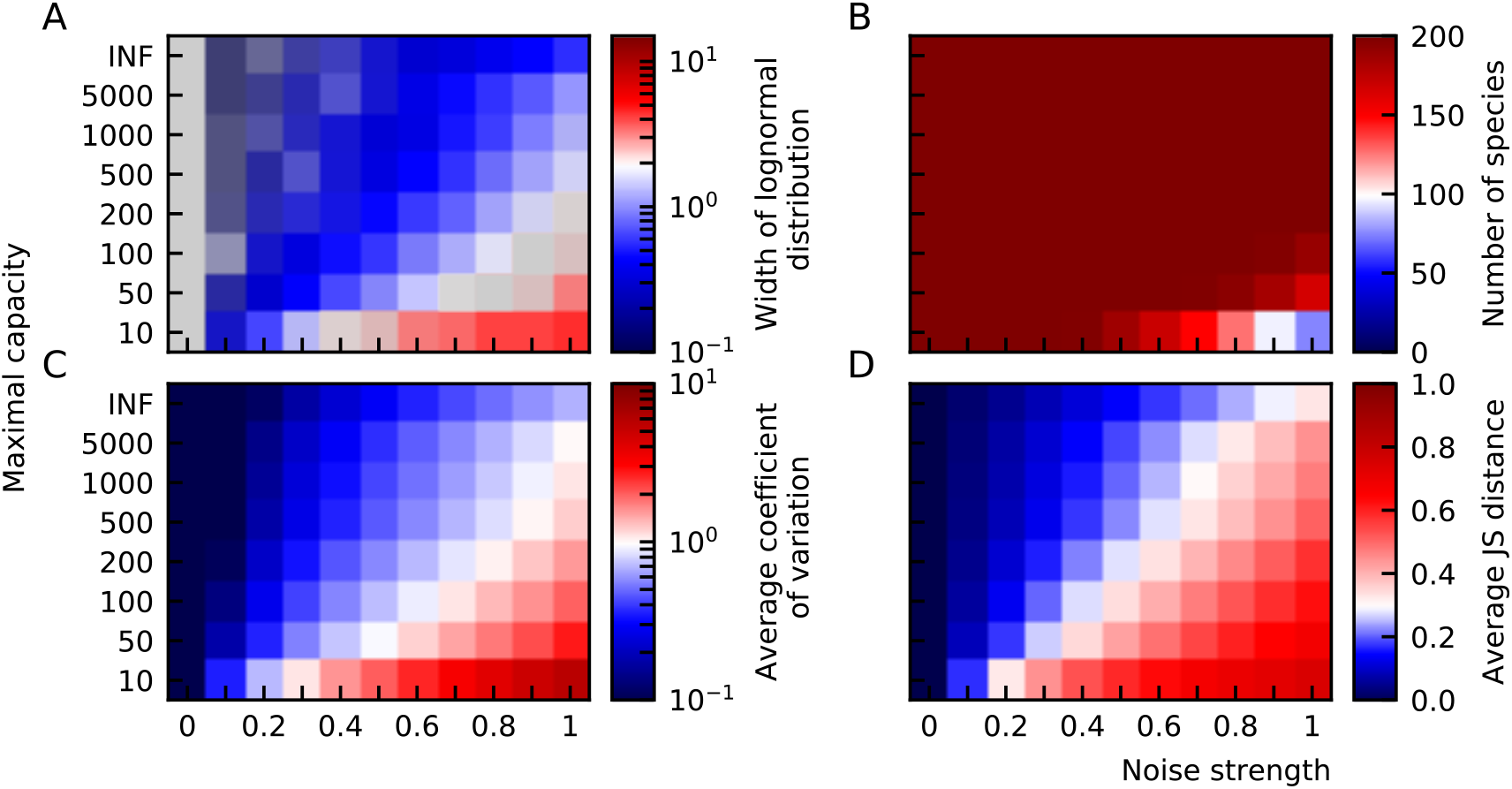
Heavy-tailed distributions for logistic equations, the influence of the maximal capacity and noise strength. (A) Abundances mostly fit a lognormal distribution best, with a tendency to normal distributions for small noise. This transition is gradual as the width of the lognormal distribution decreases in this direction (B). The species diversity is maintained except for small maximal capacities and high noise (C). The average coefficient of variation (E) and average JS distance (F), increases for decreasing maximal capacities and increasing levels of noise. Because logistic equations are used for these plots, the fixed parameters are the interaction strength *a* = 0, the connectance *C* = 0 and immigration rate *λ* =0.

In sgLV models, we can impose the strength of the noise. In IBMs, the level of stochasticity is determined by the maximum number of individuals. For decreasing maximum number of individuals the fluctuations increase (Supporting results Figure 6. C-D). Similarly, for sgLV models, the parameters of the fluctuations increase with increasing noise and decreasing maximal capacity (Figure 5).

#### Interactions make the lognormal abundance distribution wider

We previously showed that logistic equations with a maximal capacity simply reduce to gLV equations. When including interactions (Equation 3), higher-order interaction terms remain:

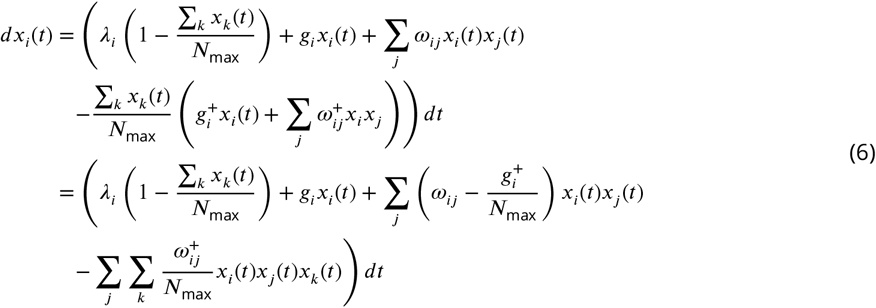

Such higher-order terms already appear in some models and are often interpreted in a phenomenological way and not as distinct mechanistic processes (Letten and Stouffer, 2019). The strength of the interaction is still only dependent on two species 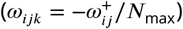 because the term originated from a pairwise interaction description.

The interaction matrix is both determined by the interaction strength and connectance. Both parameters have similar effects on the abundance distribution and fluctuations. An analysis of these parameters can be found in Supporting results.

Without maximal capacity, gLV equations are unstable for a large number of species and a large number of interactions. When a maximal capacity is imposed, the diversity does not decrease for larger connectances and interaction strengths (Supporting results Figure 10.A). This leads to lognormal distributions (Supporting results Figure 10.B). Furthermore, we see that increasing the immigration rate decreases the width of the abundance distribution (Supporting results Figure 10.C). By the implementation of a maximal capacity, we get results that are qualitatively similar to the ones of the IBM.

### Heavy-tailed distributions with logistic equations

In the absence of interactions (Equation 4), the steady-state abundance of a species *x* is given by the ratio of the growth rate to the self-interaction, −*g_i_*/*β_i_*. If the self-interaction is equal for all species—like the common convention of using *β_i_* = −1—the distribution of the steady-state abundances is obviously the distribution of the growth rates. On the other hand, if the growth rates are equal for all species, then the distribution of the steady-state abundances is the inverse distribution of the self-interactions.

#### Uniformly distributed parameters result in power law abundance distributions

If the self-interactions would be uniformly distributed between *a* and *b*, the distribution of the steady states would be

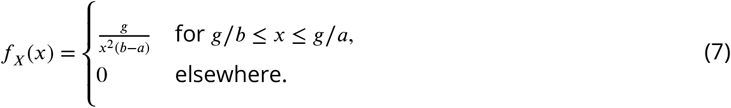

This is a power law distribution with power −2 on the bounded region [*g/b,g/a*]. The derivation of this PDF can be found in Derivations of inverses and ratio distributions.

If the distribution of growth rates and the distribution of self-interactions were uniform over [0, *g*_max_] and [-*β*_max_, 0], respectively, then the distribution of the steady states would be a ratio of uniforms distribution (RUD). By defining *ϕ* = *g*_max_/*β*_max_, we write the specific distribution of the steady states as

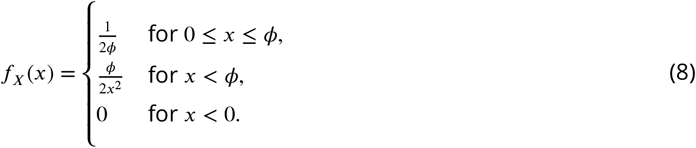

The derivation of this PDF can be found in Derivations of inverses and ratio distributions. The distribution is shown in Figure 6 and its median is *ϕ*. Experimental data corresponds well to a lognormal distribution, but can the data also be approximated by a RUD? We can calculate *ϕ* for experimental data as the median abundance value and calculate the p-value of the corresponding RUD, the results can be rejected for most of the experimental distributions (see Supporting results). It is important to keep in mind that to use this RUD, *the growth rate and self-interaction must be uncorrelated*.

**Figure 6.**
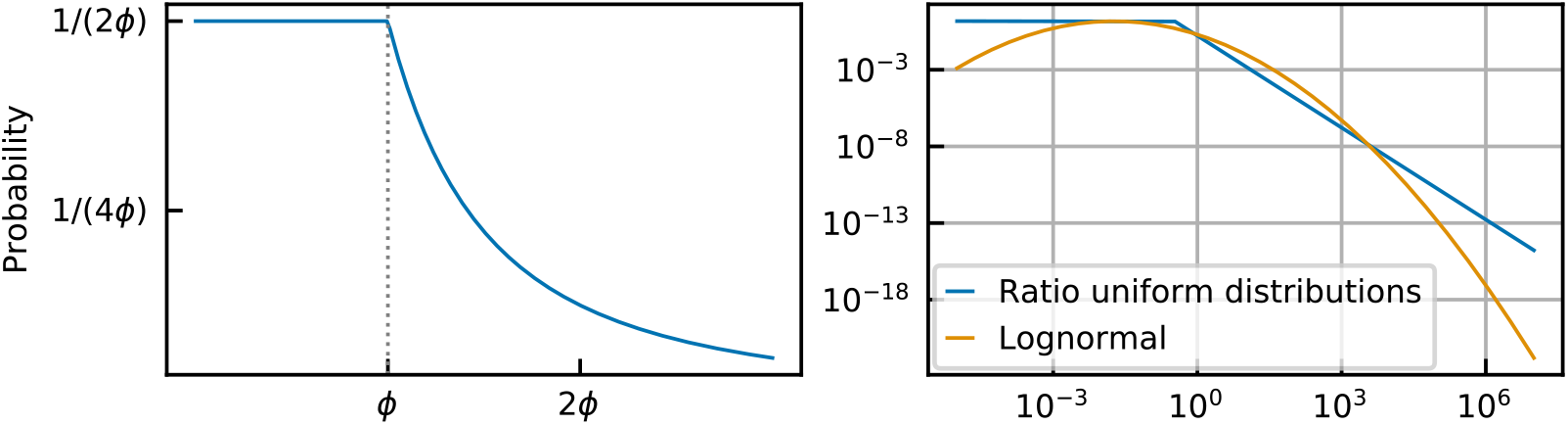
The ratio of uniforms distribution in a linear scale on the left and in a log-log scale on the right. In the right plot a lognormal distribution is drawn for comparison.

#### Lognormal abundance distributions when parameters follow lognormal distributions

If the distributions of the growth rates, and self-interactions follow a lognormal distribution, 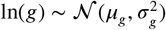 and 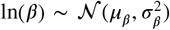, it follows that the ratio of the growth rate to the self-interaction is also a lognormal distribution,

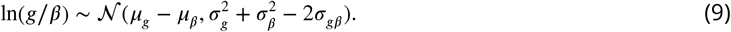

Notice that this also holds when both distributions are not independent. The derivation of this PDF can be found in Derivations of inverses and ratio distributions.

Biological growth rates and self-interactions are empiric parameters to which many microscopic processes contribute. Their effects are often multiplicative, and where multiplicative effects reign, lognormal distributions appear (Limpert et al., 2001; Zhang and Popp, 1994). Moreover, the distribution of doubling times of bacteria in the wild has been estimated to be lognormal (Gibson et al., 2018). For abundances much smaller than the carrying capacity, the growth is exponential and the doubling time is inversely proportional to the growth rate. Because the inverse distribution of a lognormal distribution is again a lognormal distribution, this implies that the bacterial growth rates follow a lognormal distribution. Furthermore, lognormal distributions of the self-interactions are not incompatible with the results of the noise color in Descheemaeker and de Buyl (2020) (see Supporting results).

From Equation 9, we conclude that the width of the abundance distribution depends on the widths of the growth rate *σ_s_* and self-interaction *σ_β_*. Fixing the width of the abundance distribution *s*, we can find the width of the growth rate given the width of the self-interaction, 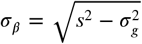 where we assumed that the distributions of the self-interaction and growth rate are independent. In this way, we can scan over different proportions of both parameters. In Figure 7, we scan over the width of the growth rates and the strength of the noise. We choose the width of the abundance *s* = 2 and, as expected, the width of the distribution remains around this value (Figure 7A). For large noise, and a large width of the growth rate distribution, a number of species goes extinct and diversity is lost (Figure 7B). In the analysis above where the growth rate and self-interactions were identical for all species, the two parameters describing the fluctuations—average coefficient of variation and average JS distance—were positively correlated. Here, with the growth rate and the self-interactions following a lognormal distribution, the parameters of fluctuations are negatively correlated with respect to the proportion of variability in the self-interaction and growth rates. For increasing widths of the growth rate and, therefore, self-interactions that become more similar for all species, the coefficient of variation increases, but the average JS distance decreases. The effect of the noise is the same as for the gLV case, it increases the parameters of the fluctuations.

**Figure 7.**
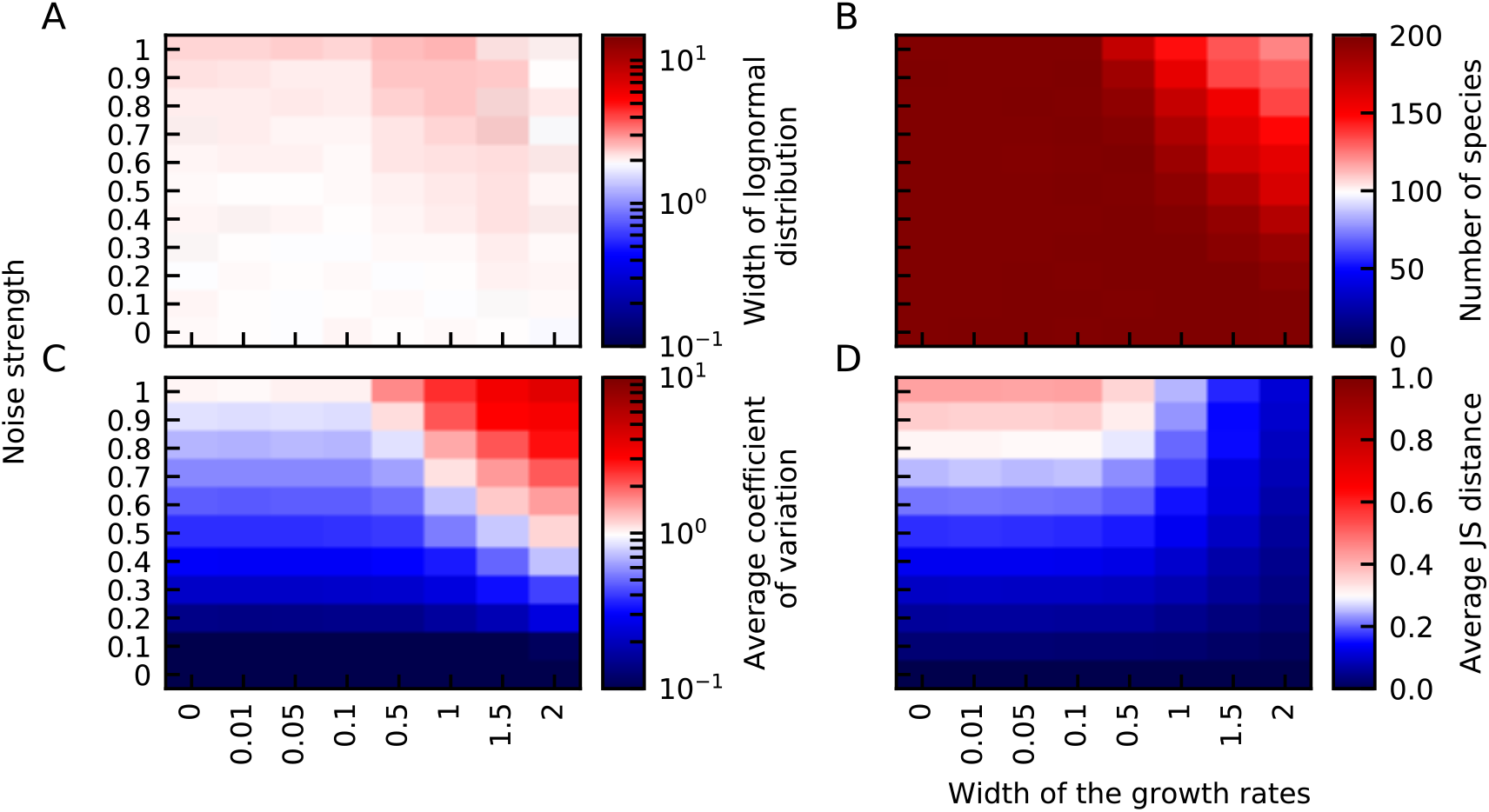
Lognormal parameter distributions lead to lognormal abundance distributions. (A) The width of the abundance distribution *s* does not depend on the width of the growth rate *σ_g_*. (B) For large noise and large width of the growth rate, species go extinct. The parameters of the fluctuation, average coefficient of variation (C) and average JS distance (D) are negatively correlated with respect to the width of the growth rate distribution. They both increase for increasing noise. The fixed parameters of these plots are the imposed width of the abundance distribution *s* = 2, the immigration *λ* =0, the interaction strength *a* = 0 and connectance *C* = 0.

## Discussion

The abundance distribution of microbial communities is heavy-tailed and typically fits a lognormal distribution. We show that lognormal abundance distributions can be obtained by different modeling approaches. In IBMs, broad distributions are the result of the system being self-organized at boundary of instability (Solé et al., 2002). Immigration increases the number of species, and interactions between species cause individuals to die. The resulting abundance distribution is lognormal-like. The width of the latter is determined by the interactions and immigration rate: larger interactions—interaction strength and connectance—and smaller immigration rates result in larger widths of the abundance distribution. In gLV models, strong interactions lead to unstable solutions and many extinct species. We consider a maximal capacity in the gLV model that mimics the maximum number of individuals in IBMs. We show that with this maximal capacity lognormal abundance distributions are obtained with increasing width for increasing connectance and interaction strength.

In the IBM and gLV model with maximal capacity, the growth rate was equal for all species and likewise for the self-interaction. The heavy-tailed rank abundance distributions were the result of interactions between species and stochasticity. Heavy tail abundance distribution can also be obtained by the appropriately setting species differences. We have shown that for logistic equations, a uniform distribution of the self-interaction leads to a power law abundance distribution, and, more iinterestingly, lognormal distributions of the growth rate and the self-interactions give rise to lognormal abundance distributions. This is in line with our expectations of such distributions.

The growth rates and carrying capactities of bacteria in the wild are unknown. In most of the cases they have not been measured as most of the species cannot be cultured in the lab. Moreover growth rates or self-interaction depend on the environment and are intrinsic properties of the species. They indeed depend on both biotic and abiotic conditions. *In vivo* experiments are even more challenging, but data of other microbes is consistent with lognormal distributed parameters (Gibson et al., 2018). The width of the distribution of growth rates and the width of the distribution of self-interactions contribute equally to the width of the final abundance distribution. However, the fluctuations in the presence of noise have a different character depending on which distribution is widest.

We expect that the heavy-tailed abundance distribution of experimental data is the result of both parameter distributions and the combined effect of interactions and stochasticity. Understanding how two disentangle these different sources of variation is essential to clarify the role of different ecological mechanisms in shaping microbial communities.

## Acknowledgments

SdB is thankful to Pierre de Buyl for helpful discussions.

## Additional files

### Supplemental information

Sources experimental data Heavy-tailed distributions

Rank abundance distribution

Kolmogorov-Smirnov test

Pearson correlation coefficient

Fano factor

Brownian motion and Ito calculus

Dissimilarity measures

Individual-based models

Fit distributions

Derivations of inverses and ratio distributions

Supplementary figures

Supporting results

### Code

All python codes to perform time series simulations, analysis and make all different figures of the main paper and supplement are available at https://github.com/lanadescheemaeker/rank_abundance.

## Notes

### Competing Interest Statement

The authors have declared no competing interest.

