## Supplemental information for "Heavy-tailed abundance distributions from stochastic Lotka-Volterra models"

**Rank abundance distribution** **5**

**Kolmogorov-Smirnov test** **6**

**Pearson correlation coefficient** **6**

**Fano factor** **6**

**Brownian motion and Ito calculus** **6**

**Dissimilarity measures** **7**

**Supplementary figures** **12**

**Supporting results** 15
The JS distance between cross-sectional data ranges from 0.4 to 0.5. 15
The width of the lognormal distribution depends on the maximal number of individuals in IBMs. 15
In the absence of interactions and immigration, maximal capacity rescales the steady state. 15
Intra-competition must be stronger than inter-competition for stability 15
Smaller maximal capacity leads to slower dynamics. 16
Interaction strength and connectance both make abundance distribution wider. 17
Experimental data does not fit the ratio of uniform distributions. 17
Self-interactions derived from noise color are compatible with lognormal distribution. 20
Diversity-complexity relationship 20
Complexity-stability in IBMs 21
Complexity-stability in PLMs 22

#### Sources experimental data

We used time series of different microbial communities. References for all of the time series can be found in Table 1.

**Table 1.** References for all time series and compositions of microbial communities.

| Label | Label in Figure 1 of the main article | Source |
|---|---|---|
| Stool A | Microbiome stool | Subject A gut of David et al. (2014) |
| Stool B | | Subject B gut of David et al. (2014) |
| Plankton bacteria | | Bacteria relative abundance of Martin-Platero et al. (2018) |
| Plankton eukarya | Plankton eukarya | Eukaryota relative abundance of Martin-Platero et al. (2018) |
| Female feces | | Feces microbiome of subject F4 at the genus level (L6) Caporaso et al. (2011) |
| Male feces | | Feces microbiome of subject M3 at the genus level Caporaso et al. (2011) |
| Female left palm | Microbiome palm | Left palm microbiome of subject F4 at the genus level (L6) Caporaso et al. (2011) |
| Male left palm | | Left palm microbiome of subject M3 at the genus level (L6) Caporaso et al. (2011) |
| Female right palm | | Right palm microbiome of subject F4 at the genus level (L6) Caporaso et al. (2011) |
| Male right palm | | Right palm microbiome of subject M3 at the genus level (L6) Caporaso et al. (2011) |
| Female tongue | Microbiome tongue | Tongue microbiome of subject F4 at the genus level (L6) Caporaso et al. (2011) |
| Male tongue | | Tongue microbiome of subject M3 at the genus level (L6) Caporaso et al. (2011) |
| Sanger | | Feces composition of DA-AD-1 of Arumugam et al. (2011) |
| Illumina | | Feces composition of MH0001 of Arumugam et al. (2011) (original data Qin et al. (2010)) |
| Pyroseq | | Feces composition of TS1_V2_turnbaugh of Arumugam et al. (2011) (original data Turnbaugh et al. (2009)) |

#### Heavy-tailed distributions

There is the misconception that most variables follow a Gaussian distribution. We even call it the "normal" distribution. The *central limit theorem* states that sums of independent random variables tend towards Gaussian distributions as the number of terms increases, even when these individual random variables do not follow a Gaussian distribution. Although this theorem is very useful and the normal distribution is omnipresent, there are as many natural processes that do not combine variables through sums, but rather through products, extremum functions, or other more complicated nonlinear functions. In that case, we observe the emergence of heavy-tailed distributions rather than normal distributions. Such distributions are ubiquitous. In this section, we first define the concept of heavy-tailed distributions mathematically. Next, we present three of the most common heavy-tailed distributions and some of their applications.

Heavy-tailed distributions are defined as distributions with tails that are heavier than tails of exponential distributions. We can define them by considering their cumulative distribution function $F$ (CDF) and more specifically the complementary cumulative distribution function $1 - F$ (Nair et al., 2020)

A distribution is called heavy-tailed if and only if for its cumulative distribution function $F$, for all $\mu > 0$,

$$\lim_{x\to\infty} \sup \frac{1 - F(x)}{e^{-\mu x}} = \infty.$$

This means that there exists a threshold value for which all higher values have a larger probability than an exponential distribution. The tails of heavy-tailed distributions are said to decay more slowly than the distribution of an exponential function.

The opposite of heavy-tailed distributions are light-tailed distributions. The intuitive conceptions we have about both types of distributions are quite different. This is manifested by the catastrophe and conspiracy principles. A probability density function (PDF) $f$ defined over the positive real numbers satisfies the *catastrophe principle* if, for $X_1, ..., X_n$, independent random variables with distribution $f$ and $n > 1$,

$$\lim_{t\to\infty} P(\max(X_1, ..., X_n) > t \mid X_1 + ... + X_n > t) = 1.$$

In other words, if the sum of random variables is large, it is because one of the samples is large and not because multiple samples are large. The heavy-tailed distributions we present here (lognormal and power law) satisfy this principle.

In contrast, light-tailed distributions tend to follow the conspiracy principle. A PDF $f$ defined over the positive real numbers satisfies the *conspiracy principle* if, for $X_1, ..., X_n$, independent random variables with distribution $f$ and $n > 1$,

$$\lim_{t\to\infty} \frac{P(\max(X_1, ..., X_n) > t)}{P(X_1 + ... + X_n > t)} = 0.$$

In other words, if the sum of the random variables is larger than its expectation value, it is most probably because many individual samples had a value higher than the expectation value. Common light-tailed distributions—the Gaussian and exponential distribution—satisfy this principle.

#### Exponential distribution

The exponential distribution serves as a boundary to distinguish light- from heavy-tailed distributions. The random variable of the exponential distribution describes the time between two events in a Poisson process. Its PDF is

$$f_{\text{exp}}(x, \lambda) = \begin{cases} \lambda e^{-\lambda x} & \text{if } 0 \geq x, \\ 0 & x < 0, \end{cases} \tag{1}$$

where $\lambda$ is the rate parameter. The mean of this distribution is $\lambda^{-1}$ and the variance is $\lambda^{-2}$. The distribution describes time-intervals of Poisson processes, such as decay times of radioactive particles, but also length-intervals for processes with a constant probability per unit length (Jorgensen, 1987), such as the distance between mutations on the DNA (Kendal, 2003).

#### Lognormal distribution

A lognormal distribution, is a continuous PDF and the logarithm of its random variable is normally distributed. Many natural phenomena are characterized by a lognormal distribution. These distributions often occur for values that cannot be negative with low mean values and high variance (Limpert et al., 2001). Growth processes are usually linear multiplicative and can be described by percentage changes of the variable. An accumulation of many of such changes is multiplicative in a linear scale, but becomes linear in a logarithmic scale. The *central limit theorem* tells us that normal distributions arise by sums of random variables. Multiplication of random variables results in lognormal distributions.

The lognormal PDF is

$$f_{\text{log}}(x, \lambda) = \begin{cases} \dfrac{1}{sx\sqrt{2\pi}} \exp\left(\dfrac{-\ln^2(x)}{2s^2}\right) & \text{if } 0 < x, \\ 0 & \text{otherwise,} \end{cases} \tag{2}$$

The theorem introduced in the previous paragraph, which states that the multiplication of random variables is a lognormal distribution, is universal and can be applied in many fields. In econometrics, it is known as *Gibrat's law*, which stipulates that the growth rate of companies is independent of their actual size, and that as a result the distribution of company sizes is lognormal (Gibrat, 1930). Other examples of lognormal distributions in nature and social behavior, are the distribution of the firing rate of neurons (Buzsáki and Mizuseki, 2014), blood pressure (Gaddum, 1945), and the size of the length of posts on internet fora (Sobkowicz et al., 2013).

##### Pareto or power law distribution

Power laws appear in a wide variety of areas such as ecology, economics, seismology, and demography (Newman, 2005). One unique and useful characteristic of the power law function is its *scale invariance*. After function variable $x$ is rescaled, the function remains proportional to the original function:

$$f_{\text{pow}}(cx) = (cx/x_m)^{-\alpha} = c^{-\alpha} f_{\text{pow}}(x) \propto f_{\text{pow}}(x).$$

Because the characteristics of this function remain the same for any scale, they are self-similar, fractal-like. The signature of a power law is the straight line in a log-log scale with a slope equal to the exponent $\alpha$ of the power law.

A power law distribution is not bounded for arbitrary exponents and domain of the variable. The standardized power law distribution is defined for $0 \le x \le 1$ and $\alpha > -1$

$$f_{\text{pow}}(x, \lambda) = \begin{cases} \alpha x^{\alpha} & \text{if } 0 \le x \le 1, \\ 0 & \text{otherwise.} \end{cases} \tag{3}$$

To allow for exponents smaller than $-1$, the domain of the power law distribution needs to have a lower bound larger than 0. The power law distribution that is defined for $x \in \mathbb{R}_{\ge x_m}$, is also known as the *Pareto distribution* after Vilfredo Pareto who used it to describe phenomena in scientific as a well as social fields. In the context of economics, he noticed that 80% of the wealth of society belongs to 20% of the population. This observation became known as the *80-20 rule* or the *Pareto principle*. Some other of the many examples of power law distributed quantities are the sizes of craters (Neukum and Ivanov, 1994), solar flares (Lu and Hamilton, 1991), cities (Gabaix, 1999), and the length of protein sequences (Jain and Ramakumar, 1999).

The Pareto distribution is a power law PDF

$$f_{\text{pareto}}(x, \lambda) = \begin{cases} \left(\dfrac{x}{x_m}\right)^{\alpha} & \text{if } 0 < x_m \le x, \\ 0 & \text{otherwise,} \end{cases} \tag{4}$$

where $\alpha < 0$.

One can also truncate the domain by giving it both a lower and an upper bound: $f_{\text{tpow}}(x) \propto x^{\alpha}$ for $x_m < x < x_M$.

Discrete versions of this distribution are the Zipf and the zeta distributions. The Zipf distribution is discussed in the next section about the rank abundance distribution.

#### Rank abundance distribution

Next to abundance distributions, rank abundance curves are much used in population studies. However, one must be careful not to confound both. In rank abundance plots, the rank of all abundances is determined by ordering them from high to low. Such plots can have a power law shape. This was for example observed by George Zipf for the frequency of words in a language—the frequency of a word is inversely proportional to the rank of the word. This empirical characteristic is known as *Zipf's law*.

Power law rank abundance distributions are associated with power law PDFs. Here follows a short demonstration (Adamic, 2000). If the rank abundance curve follows a power law, the expected value $E[x_r]$ of the variable $x$ at rank $r$ is

$$E[x_r] = c_1 r^{\beta} \tag{5}$$

This means that there are $r$ variables that have at least this value:

$$P[x \ge c_1 r^{\beta}] = c_2 r. \tag{6}$$

After changing variables, $y = c_1 r^{\beta}$, we obtain the expression of the CDF $F$,

$$F = P[x \geq y] = \frac{c_2}{c_1^{1/\beta}} y^{(1/\beta)}. \tag{7}$$

The PDF $f$ is recovered by taking the derivative of the CDF $F$

$$f = \frac{c_2}{\beta c_1^{1/\beta}} y^{(1/\beta - 1)} \propto y^{\alpha}, \tag{8}$$

the PDF is thus power law with a coefficient $\alpha = 1/\beta - 1$.

One must be careful when drawing conclusions, because the rank abundance distribution is sometimes erroneously represented in a log-linear scale instead of a log-log scale (Wierman, 2020).

#### Kolmogorov-Smirnov test

The *Kolmogorov-Smirnov test* can be used to compare either two samples and asses how likely it is that they share an underlying distribution or it can compare a sample and a reference PDF directly and tell how likely it is that the underlying distribution of the sample is the reference distribution.

The test compares the two cumulative distribution functions $F_i$ and $F_j$ of the samples or distributions. The test statistic $D$ is defined as the maximal distance between the two curves,

$$D_{ij} = \sup_x \left| F_i(x) - F_j(x) \right|. \tag{9}$$

Using the Kolmogorov distribution, one can attribute a p-value to the statistic (Marsaglia et al., 2003), or equivalently to the hypothesis that the compared distributions are the same.

#### Pearson correlation coefficient

The Pearson correlation coefficient, also referred to as Pearson's r, is a statistic that measures the linear correlation between two variables $X$ and $Y$. It is defined as the ratio of the covariance between both variables over the product of their standard deviations $\sigma_X$ and $\sigma_Y$,

$$\rho_{X,Y} = \frac{\text{cov}(X,Y)}{\sigma_X \sigma_Y}. \tag{10}$$

The value ranges between -1 and 1, with -1 and 1 denoting an exact linear relationship with negative or positive correlation respectively. A value of 0 means no correlation.

#### Fano factor

The *Fano factor*, named after Ugo Fano, measures the dispersion of a PDF. It is defined as the ratio of the variance $\sigma^2$ to the mean $\mu$,

$$F = \frac{\sigma^2}{\mu}. \tag{11}$$

It can be interpreted as a noise-to-signal ratio. It is used in neuroscience, to characterize the variability in neural spikes.

#### Brownian motion and Ito calculus

A *Brownian motion* or *Wiener process* is described by a probability distribution over the set of continuous functions $B : \mathbb{R}_{\geq 0} \to \mathbb{R}$ which is defined by three characteristics:

1. $P(B(0) = 0) = 1$, the motion starts at the origin,
2. the motion is stationary: $\forall 0 \leq s \leq t : B(t) - B(s) \sim \mathcal{N}(0, t - s)$,
3. the increments are independent: if intervals $[s_i, t_i]$ are not overlapping than $B(t_i) - B(s_i)$ are independent.

Some properties of Brownian motion are (Lee, 2013)

- it will cross $y = 0$ infinitely often,

- it does not deviate considerably from $y = \sqrt{t}$,
- it is not differentiable anywhere and
- it is self-similar: the shape of the Brownian motion does not depend on the scale.

An important feature of such a motion is the quadratic variation, which states that the expectation value of $dW^2$ is $dt$. In a regular derivative, the square and other higher order terms of the infinitesimal $dt$ of the Taylor expansion are ignored. Due to the quadratic variation, the square term of $dW$ cannot be ignored and Ito's lemma states that for a stochastic process $X_t$:

$$dX_t = \mu dt + \sigma dW_t, \tag{12}$$

and for $f$ a smooth function, we have that

$$df(t, X_t) = \left( \frac{\partial f}{\partial t} + \mu \frac{\partial f}{\partial x} + \frac{1}{2}\sigma^2 \frac{\partial^2 f}{\partial x^2} \right) dt + \frac{\partial f}{\partial x} dW_t, \tag{13}$$

and as a consequence,

$$d(\ln x_i) = \frac{dx_i}{x_i} - \frac{(dx_i)^2}{2x_i^2}. \tag{14}$$

These integration techniques are known as *Ito calculus*.

#### Dissimilarity measures

The composition of distinct communities is different and the composition of a given community changes over time. To quantify the difference between different communities, there are multiple measures. In this section, we present the Jensen-Shannon distance which relies on the Kullback-Leibler divergence.

##### Kullback-Leibler divergence

The *Kullback-Leibler (KL) divergence* is a measure that compares two probability distributions introduced by Solomon Kullback and Richard Leibler (Kullback and Leibler, 1951). It is also known as *relative entropy* and it is used in information theory, neuroscience, machine learning, etc. In the context of community dynamics, it is defined by

$$KL(x|y) = \sum_i x_i \ln \frac{x_i}{y_i} \tag{15}$$

where $x$ and $y$ represent the relative abundance vectors of the community composition. This definition is only defined if all elements of the $x$ and $y$ vector are strictly positive. In experimental data, we nevertheless have zero values for some of the abundances. There are two possibilities for this to occur: some species are absent in a certain environment, or some species are extremely rare with respect to other species in the same environment such that their abundance is too low for the detection threshold. In order to calculate the KL divergence with zero abundance, a pseudocount ($\epsilon \lll 1$) is added to the vectors. Like the Bray-Curtis divergence, the KL divergence is not a distance, because it does not satisfy the triangle inequality. Notice also that the KL divergence is not symmetrical for $x$ and $y$. The minimal value for the KL divergence is zero, this happens when both communities have exactly the same composition. The measure is, however, not bounded above, and can become infinite for some distributions. Multiple symmetric and bounded versions of this measure exist, we introduce the Jensen-Shannon divergence in the next section.

##### Jensen-Shannon divergence and distance

The *Jensen-Shannon (JS) divergence* is a measure that compares two probability distributions which is derived from the KL divergence, but which is bounded and symmetric (Lin, 1991). It is used in bioinformatics, machine learning, social sciences, etc. In the context of community dynamics, it is defined by

$$JS(x|y) = \frac{1}{2}\left( KL\left(x\middle|\frac{x+y}{2}\right) + KL\left(y\middle|\frac{x+y}{2}\right)\right). \tag{16}$$

where $x$ and $y$ represent the relative abundance vectors of the community composition. The value of the JS divergence ranges between 0 and 1, denoting two equal community compositions and two very different compositions.The square root of the JS divergence is a metric and is also called the *JS distance*.

#### Individual-based models

The concepts speak for themselves, individual-based model models (IBM) consider individuals whereas population-level models (PLM) consider populations. Individual-based models are a useful tool when one wants to take into account the differences between individuals even when they belong to the same species. They have the set-up of an automaton, *i.e.* all individuals obey a some simple rules. Although this set of rules determines only the fate of individuals, patterns can emerge on larger scales.

##### Model of Solé

We, here, shortly present the IBM of Solé et al. (2002) (called model B in the reference) which leads to heavy-tailed distributions and the species-diversity relation. This model considers a lattice filled with a fixed number of lattice sites. Every site can either be empty or harbor one individual of a species. There is a propagule supply defined with all species present, such that individuals from any species can immigrate. With $\Sigma = \{1, 2, \ldots, S\}$ the set of all species and 0 to denote the empty site as a pseudo-species, three possible transitions are defined:

1. *Immigration*: a individual of species $A \in \Sigma$ can immigrate and replace one empty site $0$ with probability $\mu_A$
$$0 \xrightarrow{\mu_A} A$$
2. *Death*: one individual of species $A \in \Sigma$ can die or emigrate, thereby emptying a lattice site. This occurs with probability $e_A$
$$A \xrightarrow{e_A} 0$$
3. *Interaction*: An interaction matrix $\Omega_{ij}$ determines the effect of species $i$ on species $j$. For two random individuals of species $A \in \Sigma$ and $B \in \Sigma$, interaction occurs when $\Omega_{AB} > \Omega_{BA}$ with probability $\Omega_{AB} - \Omega_{BA}$. The interaction consists of an individual of $A$ replacing one of $B$
$$A + B \xrightarrow{\Omega_{AB} - \Omega_{BA}} 2A.$$

Notice that there is no growth process, no self-interaction and that the implementation of the interactions is such that there are effectively only predator-prey interactions. The equivalent interaction matrix in deterministic models would be an anti-symmetric matrix.

Because the details of the exact implementation of the model are not clear from the manuscript of Solé et al. (2002) and because growth processes, self-interaction, and interactions other than predation are absent in this model, we use a derivative of this model proposed by Heyvaert (2017)

##### Model of Heyvaert

The algorithm we here describe was developed by Heyvaert (2017) based on the model of Solé et al. (2002). We consider a system with $S$ different species and a maximal number of individuals $N$. This is equivalent to a system with a lattice with $N$ sites that can each be occupied by at most one individual. We will use this image of the lattice with species and empty sites to explain the algorithm as it is easy to understand, although the spatial aspect of the lattice is unimportant for our model and no spatial dependences are taken into account. The state of the system can be described by $\vec{x} = (x_0, x_1, x_2, ..., x_S)$ where $x_i$ represents the number of individuals of species $i$ and $x_0$ the number of empty lattice sites, equivalently we can describe the system by a lattice $X$. By the construction of the model the following constraint is satisfied at all time steps:

$$\sum_{i=0}^{S} x_i = N. \tag{17}$$

At every time step, the state vector of the system is adapted by a given number of events similar to a Monte Carlo simulation. The algorithm is repeated for a fixed number of time steps. The length of every time step is equal. The simulation rules for every time step are explained in the next paragraph.

At every time step, $N$ sites of the lattice $X(t)$ are visited at random:

- If the chosen site is empty, there is a possibility for *immigration*. One species of the species is pool is chosen and the probabilities for all species are proportional to their immigration probabilities $\mu_i$. The immigration event occurs with a probability $\mu_a$. In the case of an immigration event by species $a$, the state of the system at the subsequent time point is changed:

$$x_0 \mapsto x_0 - 1$$
$$x_a \mapsto x_a + 1$$

- If the chosen site is occupied by an individual of species $a$, a second site is chosen from the lattice at time $t$, $X(t)$.
  - *Interaction:* If the second site is an individual of species $b$, this individual has a probability of interacting with the first individual of species $a$. The probability of interaction is $|\omega_{ab}|$. For $\omega_{ab} = 0$, there is no interaction. For $\omega_{ab} \neq 0$, the sign of the interaction coefficient $\omega_{ab}$ denotes whether the abundance of species $a$ will grow or decrease:

$$\left.\begin{aligned} x_0 &\mapsto x_0 \pm 1 \\ x_a &\mapsto x_a \mp 1 \end{aligned}\right\} \text{if } \omega_{ab} \lesseqgtr 0$$

  - *Growth:* If the second site is empty, the abundance of species $a$ can grow with a probability denoted by the growth rate $r_a$:

$$x_0 \mapsto x_0 - 1$$
$$x_a \mapsto x_a + 1$$

  - *Extinction:* If the individual of first site did not die through interaction, it can still die with probability $e_a$:

$$x_0 \mapsto x_0 + 1$$
$$x_a \mapsto x_a - 1$$

To calculate the state of the system at time point $t+1$, we start from the state at time $t$ and subsequently visit $N$ times a random site of the lattice at time point $t$. Because the process is random, one site can be visited twice and another is not considered at a given time step. The events associated with this lattice site change the state at time $t + 1$. Because none of the species numbers or the number of empty sites can become negative, the events can only occur if they do not violate this assumption. The implementation of this algorithm can be found at https://github.com/lanadescheemaeker/sglv_timeseries.

#### Fit distributions

Long-tailed distributions are a subclass of *heavy-tailed distributions* and many of the common heavy-tailed distributions such as the Pareto and lognormal distribution are long-tailed (Nair et al., 2020). We are mostly interested in these distributions and we will use the terminology of heavy-tailed distributions.

##### Power law distribution

As discussed in Pareto or power law distribution, there are multiple descriptions of a power law distribution defined on different domains. Here, we fit all three distribution—power law distribution, pareto distribution, and truncated power law distribution—and choose the distribution with the best fit for the Kolmogorov-Smirnov test (see Kolmogorov-Smirnov test). The fit of the three distributions is done by the scipy.stats.powerlaw and scipy.stats.pareto of the scipy package, and the powerlaw package in Python, respectively. Performing a linear fit through the logarithm of the values using a least square estimator would be statistically incorrect. The most robust method to fit a power law is the maximal likelihood estimator (Goldstein et al., 2004). This is the method used by the python functions we cited previously.

##### Lognormal distribution

We used the scipy.stats.lognorm function of the scipy package in Python to fit lognormal distributions. Before fitting the abundance distribution, we scaled all values such that the median is 1. We then imposed $\mu = 0$ and $x_0 = 1$ in Equation 1 of the main paper.

##### Normal distribution

We used the scipy.stats.norm function of the scipy package in Python to fit normal distributions.

##### Gamma distribution

The Gamma PDF is described by

$$f_\Gamma(x) = \frac{x^{\beta-1}}{\Gamma(\beta)}\left(\frac{\beta}{\bar{x}}\right)^{\beta}\exp\left(\frac{-\beta x}{\bar{x}}\right) \tag{18}$$

where $\bar{x}$ represents the average of $x$ and $\beta = (\bar{x}/\sigma_x)^2$ is the inverse coefficient of variation squared. To fit the Gamma distributions, we used the scipy.stats.gamma function of the scipy package in Python.

#### Derivations of inverses and ratio distributions

In nature, we find variables that are the inverse or ratios of other variables. Given the probability density function (PDF) of the random variables, the PDF of the inverse or ratio can be calculated. The former is also called the *reciprocal distribution*. In this section we focus on the inverse of the uniform distribution, the ratio of uniform distributions and the ratio of lognormal distributions.

##### Inverse of uniform distribution

If $z = 1/x$, a random variable, we can determine the PDF of $z$, $f_Z$, given the distribution of $x$, $f_X$. For the derivation, we rely on the cumulative density functions (CDFs) $F_X$ and $F_Z$:

$$F_Z(z) = \Pr(Z \leq z) = \Pr\left(X \geq z^{-1}\right) = 1 - \Pr\left(X < z^{-1}\right) = 1 - F_X(z^{-1}). \tag{19}$$

Because the probability density function $f$ is the derivative of the cumulative distribution $F$, we have

$$f_Z(z) = z^{-2} f_X(z^{-1}). \tag{20}$$

Therefore, if $x$ is a random variable uniformly distributed between $a$ and $b$ ($X \propto \mathcal{U}(a, b)$), or

$$f_X(x) = \begin{cases} \frac{1}{b-a} & a \leq x \leq b, \\ 0 & \text{otherwise,} \end{cases} \tag{21}$$

we obtain

$$f_Z(z) = \begin{cases} \frac{1}{z^2(b-a)} & 1/b \leq x \leq 1/a, \\ 0 & \text{otherwise.} \end{cases} \tag{22}$$

##### Ratio of uniform distributions

Suppose we have a variable $Z = Y/X$ which is the ratio of variables $X$ and $Y$, then the probability distribution of $Z$ can be determined from the joint probability of $X$ and $Y$, $f_{X,Y}$. We will, in our derivation, assume that the distributions of $X$ and $Y$ are independent such that $f_{X,Y} = f_X \cdot f_Y$. To determine the distribution of the ratio $Z$, we use the following formula (Curtiss, 1941):

$$f_Z(z) = \int_{-\infty}^{+\infty} |x| f_X(x) f_Y(xz) dx. \tag{23}$$

With $X$ and $Y$ uniformly distributed between 0 and a and b respectively—$X \propto \mathcal{U}(0, a), Y \propto \mathcal{U}(0, b)$, we have

$$f_X(x) = \begin{cases} \frac{1}{a} & 0 \leq x \leq a, \\ 0 & \text{otherwise,} \end{cases} \tag{24}$$

and

$$f_Y(y) = \begin{cases} \frac{1}{b} & 0 \leq y \leq b, \\ 0 & \text{otherwise,} \end{cases} \tag{25}$$

and the distribution of $Z$ becomes

$$f_Z(z) = \int_0^a |x| f_X(x) f_Y(xz) dx \tag{26}$$

$$= \begin{cases} \int_0^a \frac{|x|}{ab} dx = \frac{a}{2b} & \text{for } 0 \leq z < \frac{b}{a} \\ \int_0^{b/z} \frac{|x|}{ab} dx = \frac{b}{2az^2} & \text{for } z \geq \frac{b}{a}. \end{cases} \tag{27}$$

#### Ratio and inverse of lognormal distributions

Suppose $X$ and $Y$ follow a lognormal distributed, such that $\ln(X)$ and $\ln(Y)$ describe normal distributions. The ratio distribution of these lognormal distributions $Y/X$ can also be written as $\exp(\ln(Y/X)) = \exp(\ln(Y) - \ln(X))$. If we define the mean of $\ln(X)$ as $\mu_X$ and similarly for $\ln(Y)$, and the covariance matrix between both distributions is

$$\text{cov} = \begin{bmatrix} \sigma_X^2 & \sigma_{XY} \\ \sigma_{XY} & \sigma_X^2, \end{bmatrix} \tag{28}$$

then $Z = \ln(Y) - \ln(X)$ is normally distributed with mean $\mu_Y - \mu_X$ and variance $\sigma_X^2 + \sigma_Y^2 - 2\sigma_{XY}$. Therefore, $Y/X = \exp(Z)$ follows a lognormal distribution.

Following the same logic, one obtains that the inverse distribution of a lognormal distribution is a lognormal distribution:

$$\ln(X) \propto \mathcal{N}(\mu, \sigma^2) \Leftrightarrow \ln(1/X) \propto \mathcal{N}(-\mu, \sigma^2). \tag{29}$$

### Supplementary figures

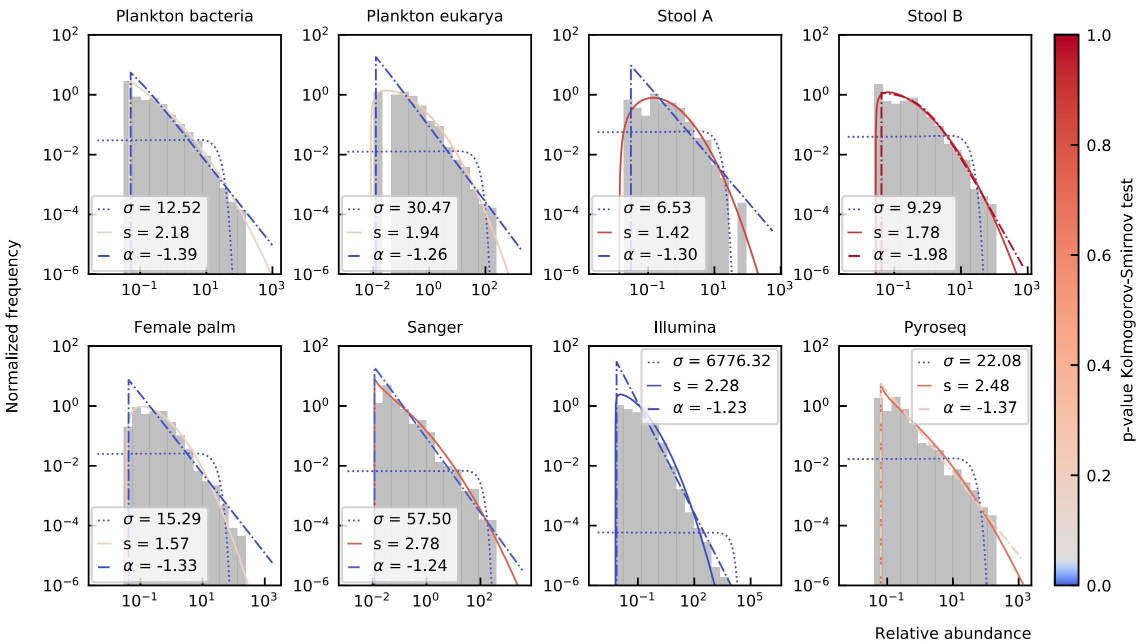

**Figure 1.** Experimental data has a heavy-tailed distribution. The lognormal distribution (full line) fits best. The power law (dash-dotted) and exponential (dotted) distribution are rejected for most communities because the p-values of the KS test are lower than 0.05 (lines are colored in blue). The widths $s$ of the lognormal distribution range between 1.42 and 2.78. For data coming from a time series, we considered the first time point.

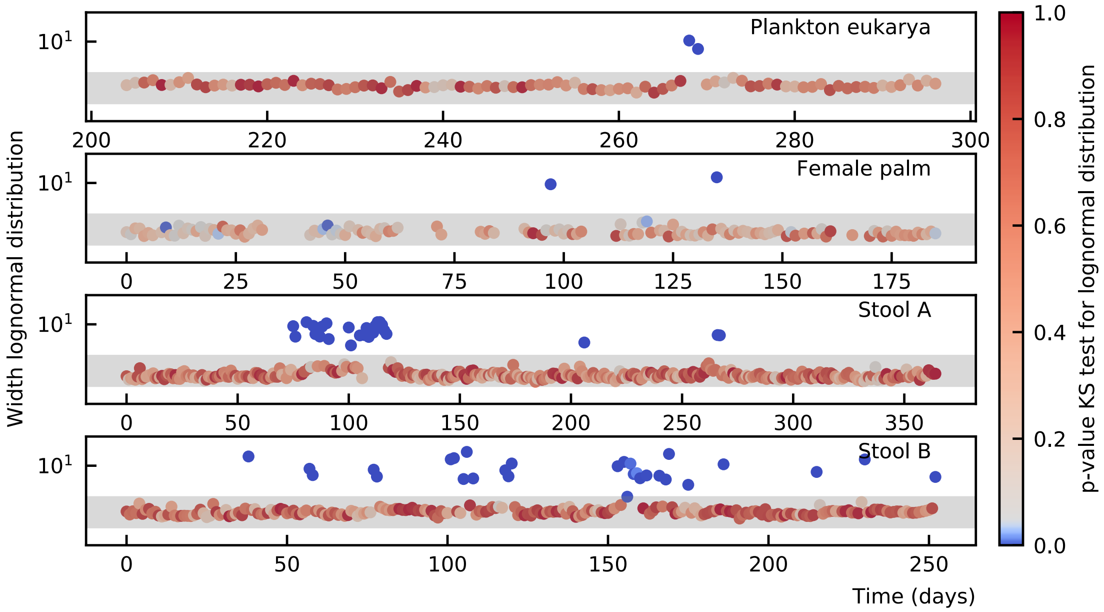

**Figure 2.** The species abundances of microbial communities show large fluctuations over time, but the width of the abundance distribution remains nearly stable over time. The grey horizontal band extends from 1 to 3. The color of the dots shows the p-value of the KS test to a lognormal distributions, blue values are poor fits. The lognormal fits for which the width is higher than 3, can all be rejected.

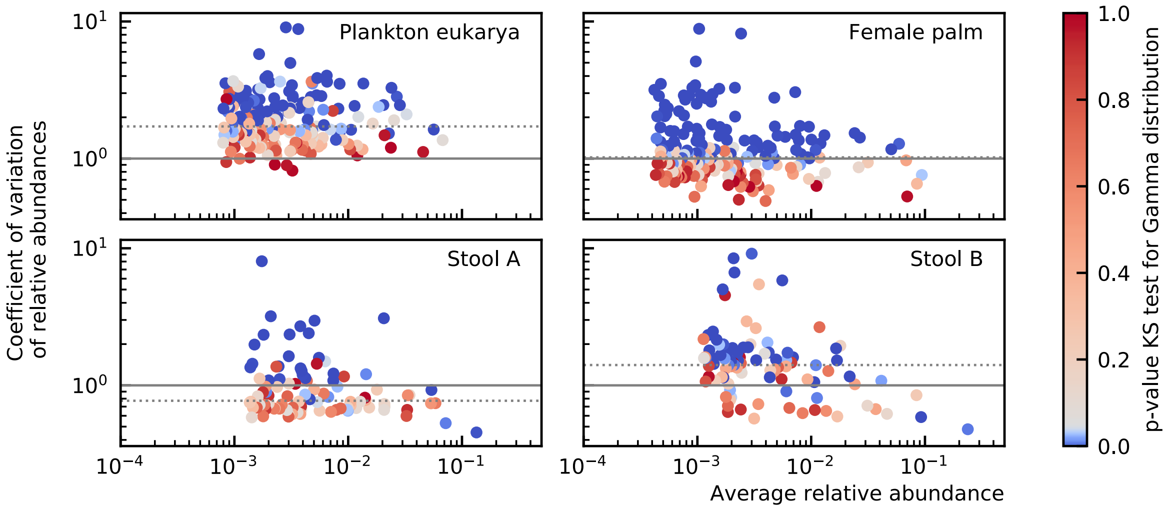

**Figure 3.** Longitudinal experimental data of microbial communities has a coefficient of variation that is around one. For species with a large coefficient of variation, we can reject the hypothesis that the population follows a Gamma distribution. The dotted horizontal lines denote the median coefficient of variation per community.

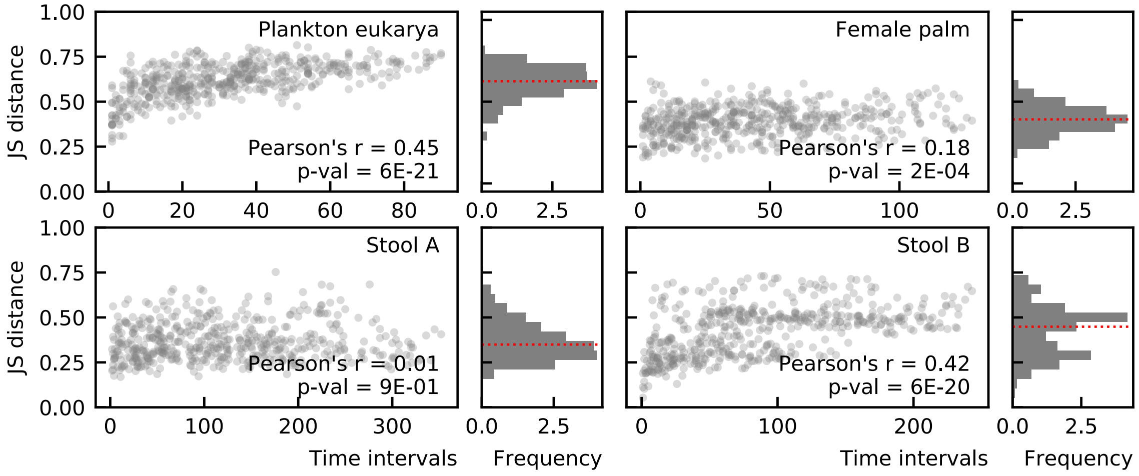

**Figure 4.** Jensen-Shannon (JS) distance of longitudinal experimental data as a function of the time interval between both compositions. The median value (red line) ranges from 0.3 to 0.4. The Pearson correlation function and p-value for non-correlation are given.

#### Supporting results

##### The JS distance between cross-sectional data ranges from 0.4 to 0.5.

We compare the gut microbial composition of different individuals using the Sanger, Illumina, and Pyroseq datasets (see Table 1 for the references). The median JS distance between compositions of distinct individuals is between 0.4 and 0.5 (Figure 5).

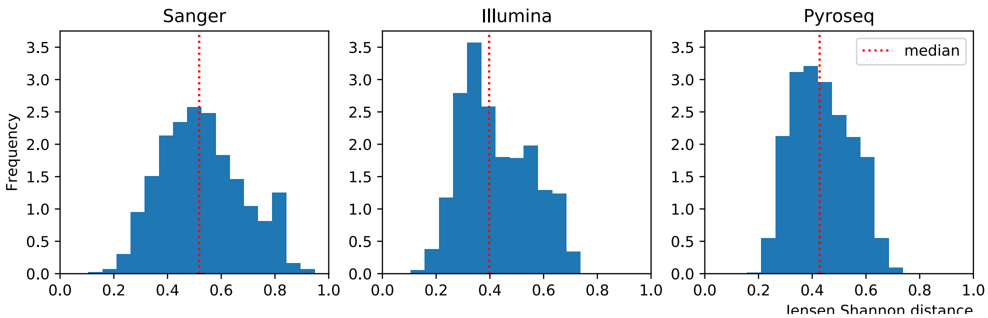

**Figure 5.** Jensen-Shannon distances between different individuals of datasets Sanger, Illumina and Pyroseq. The median JS distance is 0.4 to 0.5.

##### The width of the lognormal distribution depends on the maximal number of individuals in IBMs.

The width of the lognormal distribution increases for smaller maximum numbers of individuals and immigration (Figure 6). As the width of the lognormal increases, the number of species decreases. Simultaneously, the average coefficient of variation and the average JS distance increase.

##### In the absence of interactions and immigration, maximal capacity rescales the steady state.

For gLV equations (Equation 6 in the main paper) without immigration, we know that the non-trivial steady state is given by $x_i^\dagger = -\sum_j (\omega^{-1})_{ij} \cdot g_j$. This state is possibly non-feasible if the steady state of one of the species is negative. Because we know that the logistic equations with a maximal capacity reduce to gLV equations, we can calculate the steady state for logistic equations with a maximal capacity. The interaction matrix is

$$\omega_{ij} = -\frac{g_i^+}{N_{\text{max}}} - \beta_{ii}\delta_{ij} \tag{30}$$

where $\delta_{ij}$ is the Kronecker delta. The Sherman-Morrison formula allows us to invert this interaction matrix and we obtain the expression for the steady state (Sherman and Morrison, 1950)

$$x_j^* = \beta_j^{-1} g_j \left(1 + \sum_i \beta_i^{-1} g_i / N_{\text{max}}\right)^{-1}. \tag{31}$$

Because the steady state without maximal capacity is $g_i/\beta_i$, we see that the maximal capacity rescales the steady state by a factor $(1 + \sum_i x_i^\dagger / N_{\text{max}})^{-1}$. Therefore the evenness of the species abundances or the steepness of the rank abundance is not changed when a maximal capacity is added in the absence of immigration and interactions.

##### Intra-competition must be stronger than inter-competition for stability

Consider a system of gLV equations

$$\dot{x}_i = (g_i + \sum_j \omega_{ij} x_j) x_i \tag{32}$$

The Jacobian of this equation is

$$J_{ij} = x_i^* \omega_{ij} \tag{33}$$

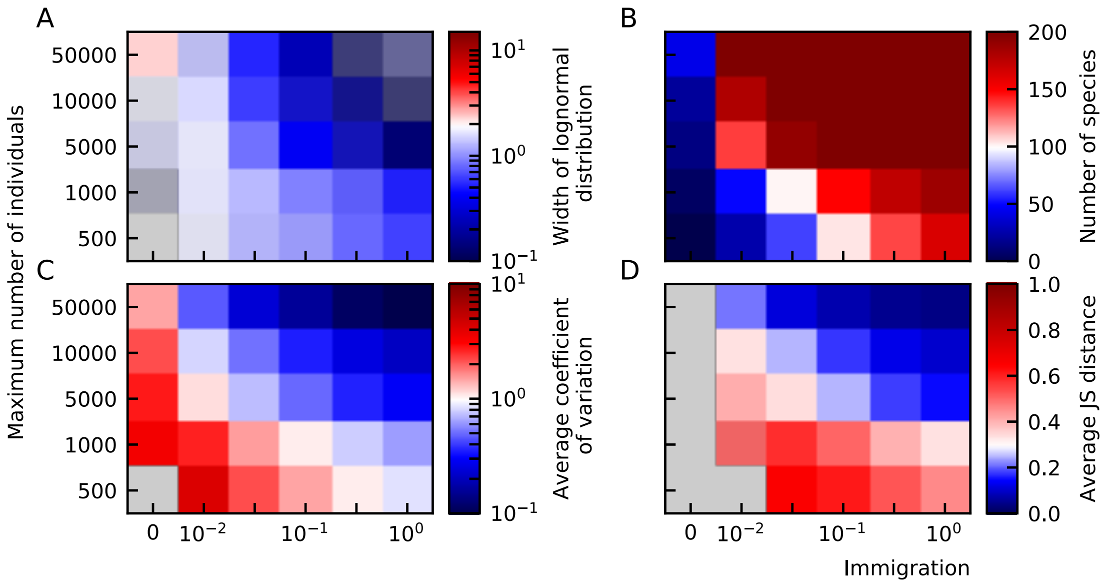

**Figure 6.** Individual-based models result in heavy-tailed abundance distributions. (A) The width of the lognormal distribution increases for smaller maximum numbers of individuals and immigration. As the width of the lognormal increases, the number of species decreases (B). Simultaneously, the average coefficient of variation (C) and the average JS distance (D) increase. The fixed parameters of these plots are the interaction strength 0.5 and the connectance 0.5.

---

where $x_i^*$ represents the steady-state value of $x_i$. For all species equivalent the Jacobian takes the shape

$$J = x^* \begin{bmatrix} d & o & \dots & o \\ o & d & \dots & o \\ \vdots & \vdots & \ddots & \vdots \\ o & o & \dots & d \end{bmatrix} \tag{34}$$

where the diagonal elements $d$ denote the self-interaction and the off-diagonal elements $o$ the interaction between species. The eigenvalues of this matrix are $x^*(d-o)$ and $x^*((N-1)o+d)$. Stability thus requires that $d < o$ for otherwise the first eigenvalue is positive and $d < o < 0$ to keep the second eigenvalue negative. When the inter-species competition is larger than the self-interaction, i.e. $d > o$ (since both values are negative), the system is unstable and a small perturbation of the steady state will result in one of the species taking over (winner-take-all). Because the Jacobian of the logistic model with maximal capacity is given by Equation 30, and we have $-\frac{g_i^+}{N_{\mathrm{max}}} - \beta_{ii} < -\frac{g_i^+}{N_{\mathrm{max}}} < 0$, the logistic model with maximal capacity is stable.

#### Smaller maximal capacity leads to slower dynamics.

The steady state of Equation 5 of the main paper is

$$x_i^* = \frac{g_i}{\beta_i} + \frac{g_i^+}{\beta_i} \frac{\sum_j g_j \beta_j^{-1}}{N_{\mathrm{max}} - \sum_j g_j^+ \beta_j^{-1}}. \tag{35}$$

When $g = g^+$, this simplifies to $x_i = y_i/(1 + \sum_j y_j/N_{\mathrm{max}})$ where $y$ is the steady-state value of $x$ in the absence of a maximal capacity $y_i = g_i\beta_i^{-1}$. The speed of the dynamics to steady state can be estimated by evaluating $\frac{\mathrm{d}\dot{x}_i}{\mathrm{d}x_i}$ in $x = x^*$. The derivative is

$$\frac{\mathrm{d}\dot{x}_i}{\mathrm{d}x_i} = g_i - \frac{g_i^+}{N_{\mathrm{max}}} \left( \sum_j x_j + x_i \right) - 2\beta_i x_i \tag{36}$$

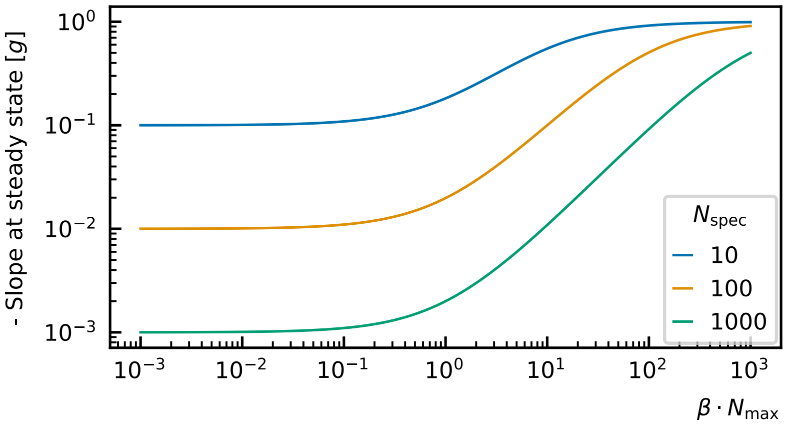

**Figure 7.** The speed with which a system goes to steady state is related to the slope of the derivative at the steady state. The smaller the maximal capacity the smaller this slope, the lower the speed. The speed is shown for systems where all species are equal ($\beta_i = \beta$, $g_i = g_i^+ = g$ for all $i$).

Evaluated this equation in the steady state, we estimate the speed to the steady state by

$$\left.\frac{\mathrm{d}\dot{x}_i}{\mathrm{d}x_i}\right|_{x^*} = \left(-g_i + g_i^+ \frac{\Sigma^- + \Sigma^+}{N_{\text{max}} + \Sigma^+}\right) \cdot \left(1 + \frac{g_i^+}{\beta_i N_{\text{max}}}\right) \tag{37}$$

with $\Sigma^{\pm} = \sum_j g_j^{\pm} \beta_j^{-1}$. For large $N_{\text{max}}$, the speed reduces to $-g_i$ which is the speed for the normal logistic equation at steady state. If we assume that all species are equivalent $\beta_i = \beta$ and $g_i = g_i^+ = g$ for all $i$ (or equivalently $g_i^- = 0$), we can estimate the speed for small $N_{\text{max}}$ by $-g/N_{\text{spec}}$ with $N_{\text{spec}}$ the number of species (Figure 7).

#### Interaction strength and connectance both make abundance distribution wider.

The interaction matrix is an important factor for the shape of the abundance distribution. Both the interaction strength and connectance strengthen each other (Figure 9). When the maximal capacity increases, the width of the lognormal distribution increases. In the absence of a maximal capacity, the solution for large interaction and connectance is unstable and many species go extinct.

#### Experimental data does not fit the ratio of uniform distributions.

For a logistic model with uniformly distributed growth rates and self-interactions, the abundances follow a ratio of uniform distributions (RUD). Its description is given by Equation 8 of the main paper where the parameter $\phi$ is the median of the distribution. Without free parameters, we can assess how likely it is that the abundances follow this distribution. We compare the RUD to a lognormal distribution, they are denoted by the full and dashed lines in Figure 11. Notice that the latter distribution needs to be fitted. We asses the likelihood for both distributions by the Kolmogorov-Smirnov test (see Kolmogorov-Smirnov test) and conclude that we can reject the hypothesis that the abundances follow a RUD for many of the experimental distributions (blue lines) and that the lognormal distribution is more likely for all experimental data but the microbiome of the female palm.

One important element is that not all reads of the sample could be classified and that a considerable percentage of the data is missing. In the Sanger, Illumina and Pyroseq data, it amounts up to 50%. When we assume that these reads come from species that are undersampled and have a lower abundance than the smallest abundance measured, we need to add a lot of small abundance species to the histogram shifting the median value to lower values and, therefore the knee of the RUD to the left. In that case, the hypothesis that the abundance distribution follows a RUD can be rejected for all experimental data considered.

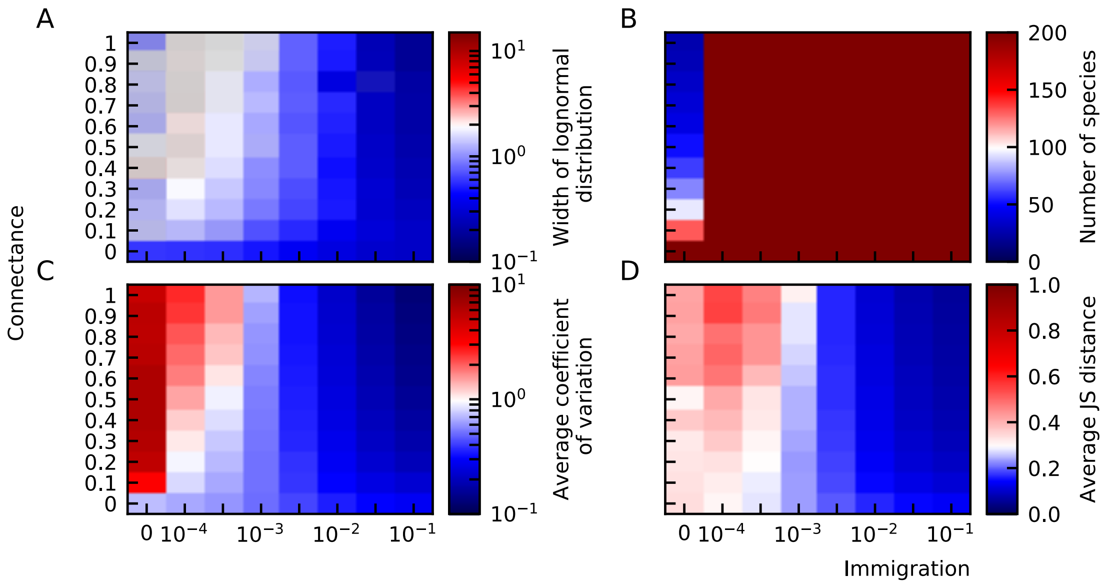

**Figure 8.** Generalized Lotka-Volterra models with a maximal capacity result in heavy-tailed abundance distributions. (A) The width of the lognormal distribution becomes larger for decreasing immigration and increasing connectance. (B) Diversity is maintained in the presence of immigration. The average coefficient of variation (C) and the average Jensen Shannon distance (D) grow for decreasing immigration. The number of species is $N_{\text{spec}} = 200$. The fixed parameters of these plots are the interaction strength $\omega = 0.5$, the noise strength $\eta = 0.5$ and the maximal capacity 100.

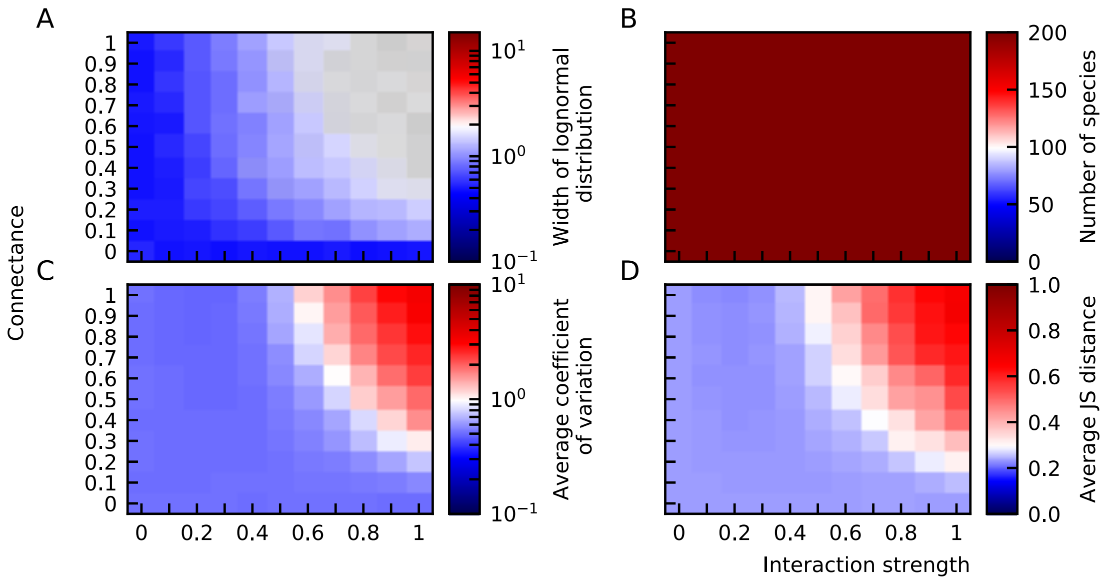

**Figure 9.** The width of the abundance distribution (A), the average coefficient of variation (C) and average JS distance (D) increase for larger interaction strength and connectance. The diversity is maximal (D). The number of species is $N_{\text{spec}} = 200$. The fixed parameters of these plots are strength of the noise $\eta = 0.5$, the immigration rate $\lambda = 0.1$ and the maximal capacity $N_{\text{max}} = 100$.

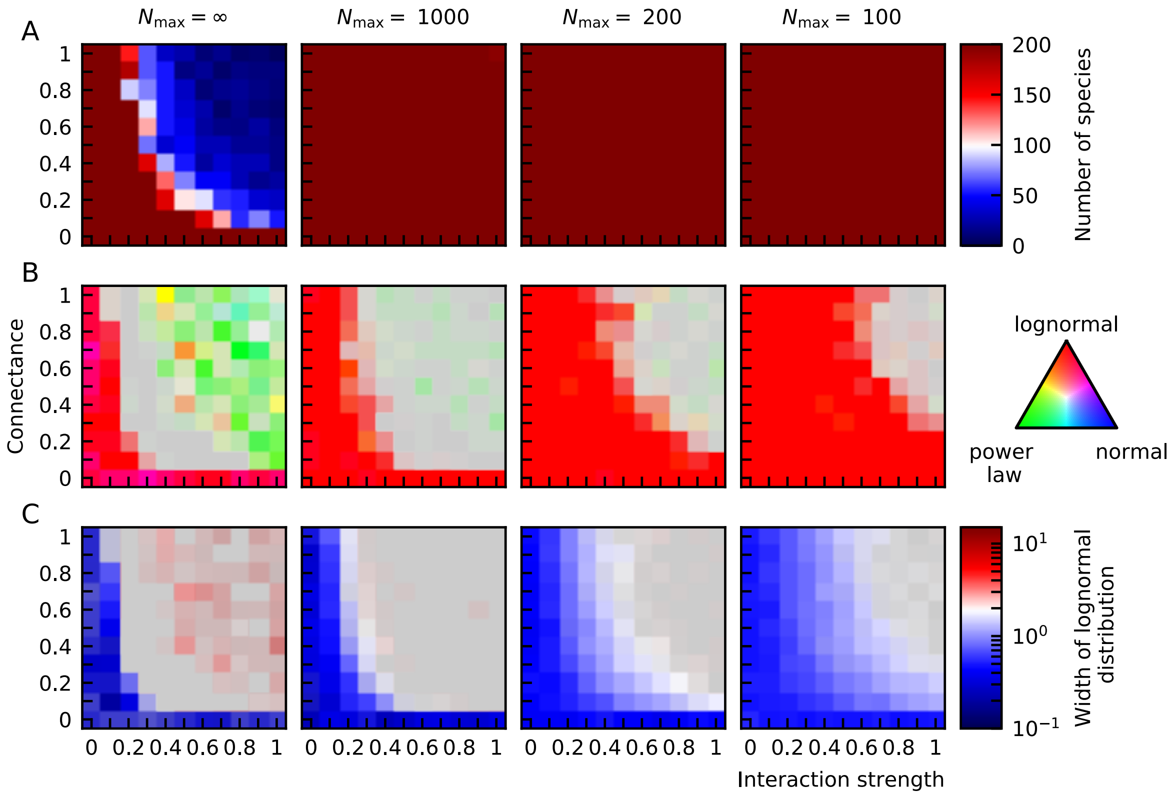

**Figure 10.** The width of the lognormal distribution as a function of the interaction strength and connectance for different values of the maximal capacity $N_{max}$. The grey areas denote unstable solutions. Smaller maximal capacities allow for stable solutions with more and larger interactions. More and larger interactions lead to wider abundance distributions.

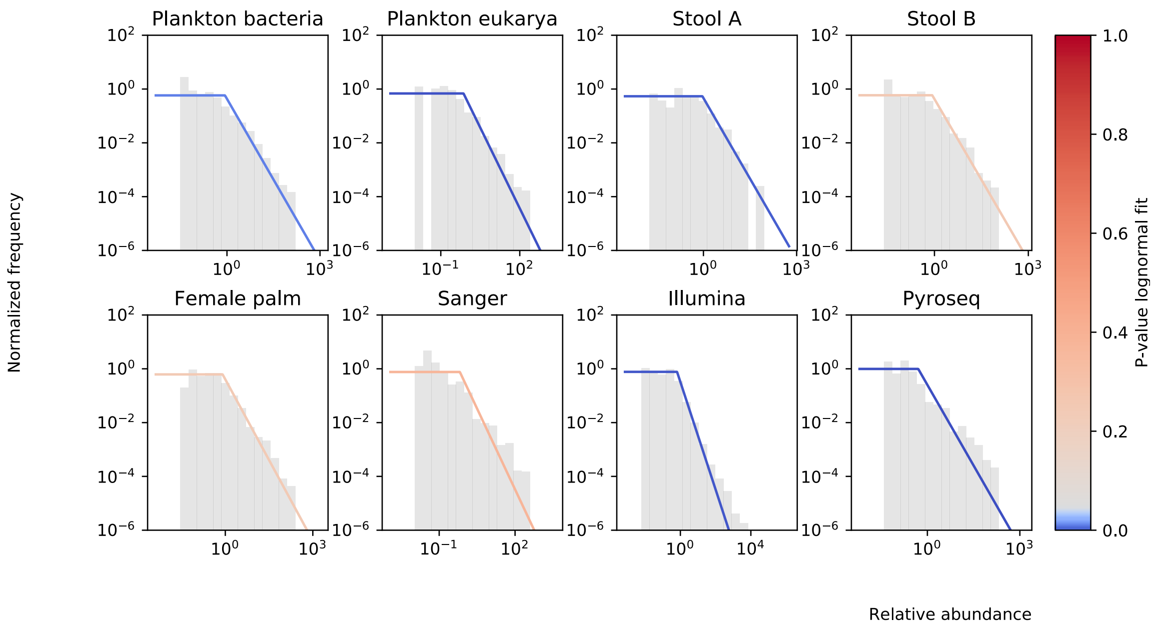

**Figure 11.** Experimental data compared to their RUD curve using $\phi$ as the median value of the measured abundances.

#### Self-interactions derived from noise color are compatible with lognormal distribution.

In Descheemaeker and de Buyl (2020), we have shown that, assuming the experimental data can be modeled by logistic equations, the self-interaction rate can be determined given the abundance and noise color, which is defined as the slope of the power spectral density (Figure 12B). We determined the self-interactions and growth rates assuming a certain sampling rate ($\delta t = 0.05$), and concluded that these results are not incompatible with a lognormal distribution for both variables. This result is not highly surprising, because the abundance distribution of the experimental data, which is lognormal, was used to infer the parameters. Nevertheless, it shows that the results of the noise color are compatible with lognormal distributions of the parameters.

#### Diversity-complexity relationship

There is a trade-off between the complexity of a network and its stability. The complexity is related to the number of species $S$, number of interactions or connectance $C$ and the strength of the interactions. Consider a community matrix—defined as the Jacobian of a system evaluated in the fixed point—with -1 on the diagonal and mean 0 and variance $\xi^2$ of the off-diagonal elements. In this case, May found that systems for which

$$\xi\sqrt{SC} < 1 \tag{38}$$

have a high probability to be stable. Using random matrix theory Allesina and Tang (2015) generalized this inequality to community matrices with random elements on the diagonal. In that case the inequality becomes

$$\xi\sqrt{SC} < \bar{d} \tag{39}$$

where $-\bar{d}$ is the average of the diagonal elements of the community matrix. Remarkably, the distribution of the off-diagonal elements in the community matrix is unimportant (Allesina and Tang, 2015) and although the normal distribution is often used, uniform distributions lead to similar results provided the appropriate variance is used and the mean is zero. Furthermore, they defined the criterion in the case that the mean of the off-diagonal elements is $\mu \neq 0$, but assuming the diagonal elements $-d$ are all identical,

$$\max\left(\sqrt{SC(\xi^2 + (1-C)\mu^2)} - C\mu, (S-1)C\mu\right) < d. \tag{40}$$

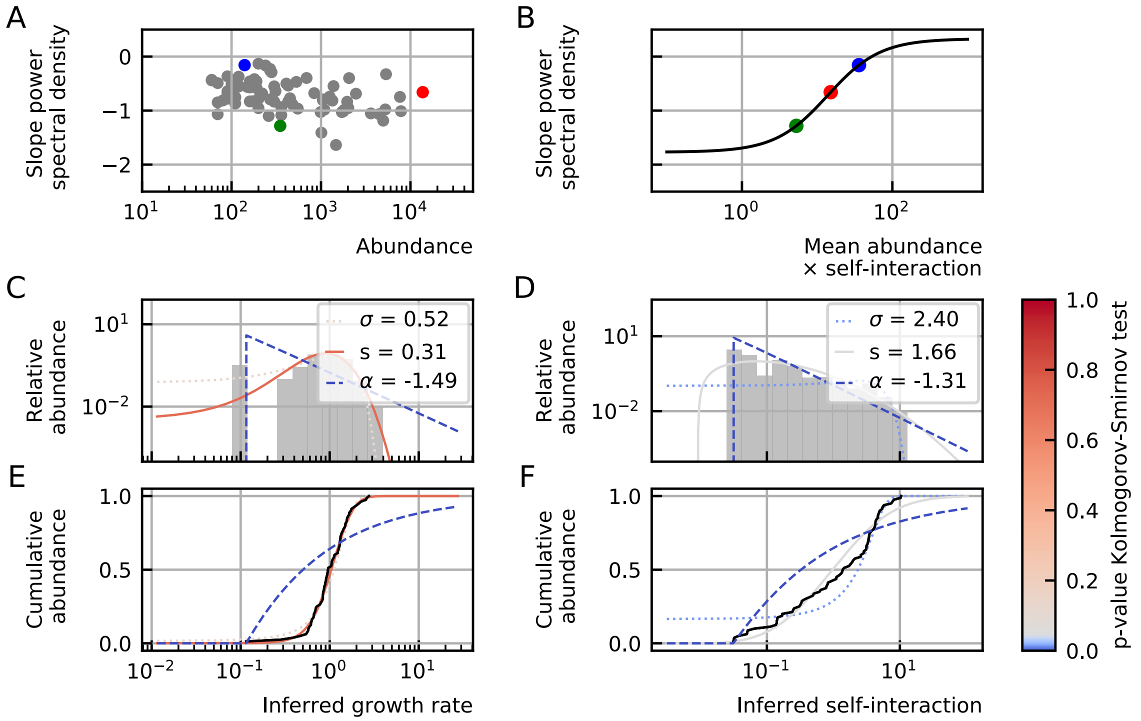

**Figure 12.** (A) The noise color as a function of the abundance for the David Stool A data. (B) The simulated relation between the noise color, mean abundance and self-interaction. (C) The self-interaction inferred from the noise color and abundance. (D) The distribution of the inferred self-interactions fitted by a normal (dotted line), lognormal (full line) and power law (dashed line). The color of the lines denotes the p-value of the Kolmogorov-Smirnov test. The lognormal curve has the best fit. (E) The cumulative density function of the self-interactions (black line) with the fitted lines corresponding to the ones in (E)

A result of May's instability criterion (Equation 38) is that the instability border is a hyperbolic relationship between the number of species, or diversity $S$ and the connectance $C$: $S \propto C^{-1}$. Without the assumption of May that the mean of the off-diagonal elements of the community matrix are zero, the relationship becomes more complicated (Equation 40).

For gLV models without immigration (Equation 2 in the main text with $\lambda_i = 0$), the community matrix is

$$M_{ij} = x_i^* \omega_{ij} \tag{41}$$

where $x_i^*$ is the steady state given by $x_i^* = \sum_j (\omega^{-1})_{ij} g_j$. From the definition of the community matrix, one sees that its connectance is equal to the one of the interaction, but that the distribution of elements of the community matrix is distorted by the abundance distribution. Also in the presence of immigration, the steady state is altered which changes the community matrix. How these effects influence the stability is still an open question (Allesina and Tang, 2015).

#### Complexity-stability in IBMs

Montoya and Solé (2003) reported experimental data that deviated from May's hyperbolic species-connectivity relation, $S \propto C^{-1+\epsilon}$ with $0 \leq \epsilon \leq 0.5$. Solé et al. (2002) shows that an IBM with immigration can explain this relationship. Solé et al. (2002) reported that the diversity—defined as the number of (non-zero) species—and connectance are connected through a power law relation. We also include the interaction strength in our definition of complexity based on the definition of May. However, we cannot compare the complexity defined here as $a^2C$ with the one of May defined as $\xi^2 C$ as $a^2$ is the variance of the interaction matrix and $\xi^2$ the variance of the community matrix. The latter depends on the abundance distribution (Equation 41). Although there is a difference between both definitions, we still see that the definition of complexity creates a boundary for stability (Figure 13). For small complexity, we see saturation because the number of species in the immigration pool is finite ($N_{\text{spec}} = 500$) as well as the maximum number of individuals ($N_{\text{max}}$). The exponent of the power law is more negative for smaller immigration rates (Figure 13). This quantitatively agrees with the mean-field model of Solé et al. (2002). We also show how the width of the abundance distribution grows with the complexity (Figure 13).

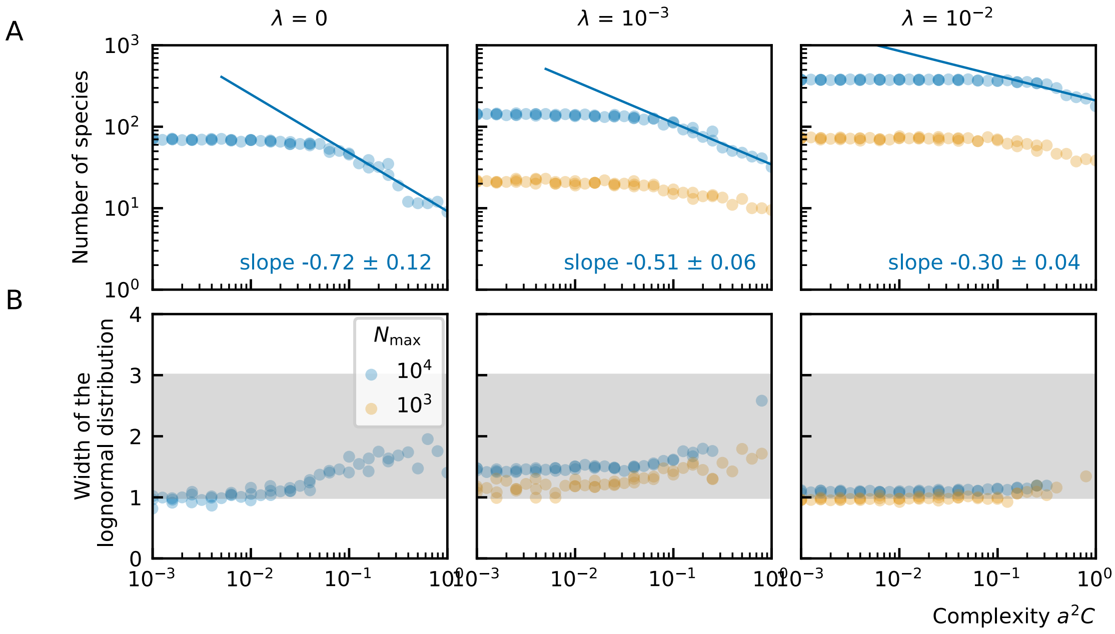

**Figure 13.** (A) Diversity-complexity relation for IBM for different immigration rates ($\lambda = 0$, $10^{-3}$ and $10^{-2}$). The solutions form a border. Because of immigration, solutions under the curve are unstable and because of interactions and a limited number of lattice sites any solutions above the curve are unstable. There is relation between the complexity and number of species can be approximated by a power law. For smaller complexity, we see saturation because the number of species in the immigration pool is finite ($N_{\text{spec}} = 500$). The exponent of the power law is more negative for smaller immigration rates. (B) The width of the abundance distribution fitted with a lognormal. The width increases for increasing complexity.

#### Complexity-stability in PLMs

For PLMs, we also analyze the relation between diversity and complexity by relying on the same definition of the complexity $a^2C$. For small complexity, there is again saturation because the number of species in the equations is finite ($N_{\text{spec}} = 500$). Similar to the results for the IBM, the slope of the power law becomes flatter for increasing immigration rates when there is a maximal capacity (Figure 14). We also show how the width of the abundance distribution grows with the complexity (Figure 13). The width of the abundance distribution increases for increasing complexity. However, for increasing complexity $a^2C > 0.05$ the hypothesis that the data follow a lognormal function is rejected (Figure 14).

As a conclusion, immigration allows for more diversity in models with a maximal capacity. The diversity-complexity relation is a power law. The width of the abundance distribution increases for increasing complexity but the distribution deviates from a lognormal distribution. Smaller maximal capacities lead to larger widths.

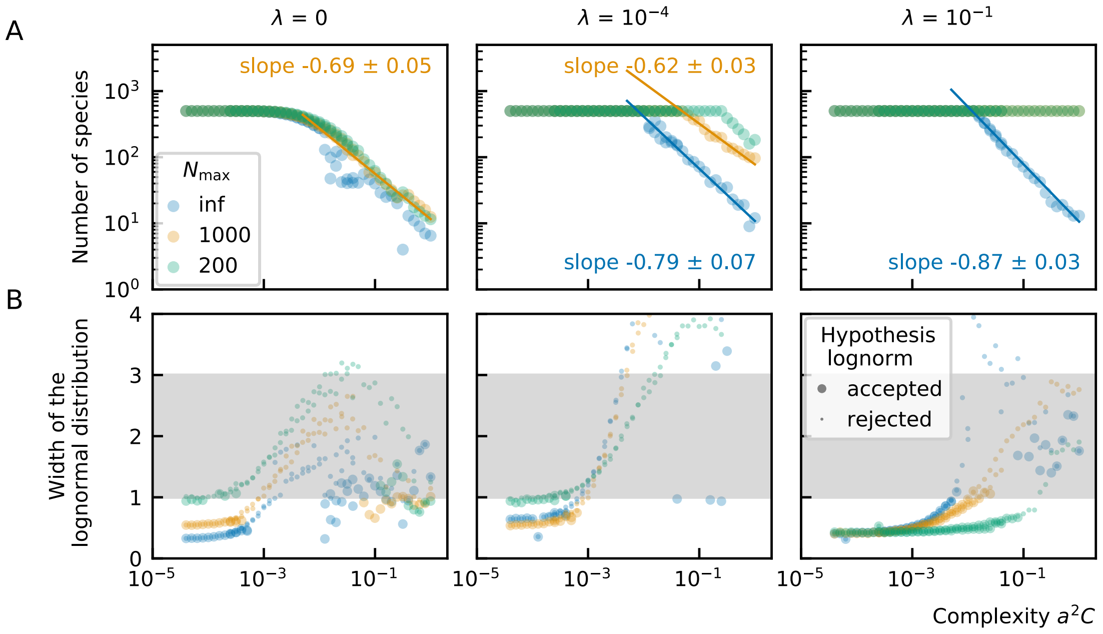

**Figure 14.** (A) Diversity-complexity relation for sgLV for different immigration rates ($\lambda = 0$, $10^{-4}$ and $10^{-1}$). There is relation between the complexity and number of species can be approximated by a power law. For smaller complexity, we see saturation because the number of species in the immigration pool is finite ($N_{\text{spec}} = 500$). The exponent of the power law is more negative for smaller immigration rates. (B) The width of the abundance distribution fitted with a lognormal. The width increases for increasing complexity, but the abundance distribution deviates from a lognormal distribution.
